# Fgf-driven Tbx protein activities directly induce *myf5* and *myod* to initiate zebrafish myogenesis

**DOI:** 10.1101/766501

**Authors:** Daniel P.S. Osborn, Kuoyu Li, Stephen J. Cutty, Andrew C. Nelson, Fiona C. Wardle, Yaniv Hinits, Simon M. Hughes

## Abstract

Skeletal muscle derives from dorsal mesoderm that is formed during vertebrate gastrulation. Fibroblast growth factor (Fgf) signalling is known to cooperate with transcription factors of the Tbx family to promote dorsal mesoderm formation, but the role of these proteins in skeletal myogenesis has been unclear. Using the zebrafish, we show that dorsally-derived Fgf signals act through Tbx16 and Tbxta to induce two populations of slow and fast trunk muscle precursors at distinct dorsoventral positions. Tbx16 binds to and directly activates the *myf5* and *myod* genes that are required for commitment to skeletal myogenesis. Tbx16 activity depends on Fgf signalling from the organiser. In contrast, Tbxta is not required for *myf5* expression. However, Tbxta binds to a specific site upstream of *myod* not bound by Tbx16, driving *myod* expression in the adaxial slow precursors dependent upon Fgf signals, thereby initiating muscle differentiation in the trunk. After gastrulation, when similar muscle cell populations in the post-anal tail are generated from the tailbud, declining Fgf signalling is less effective at initiating adaxial myogenesis, which is instead initiated by Hedgehog signalling from the notochord. Our findings provide insight into the ancestral vertebrate trunk myogenic pattern and how it was co-opted during tail evolution to generate similar muscle by new mechanisms.

## Introduction

Sarcomeric muscle arose early in animal evolution, creating a defining feature of the metazoa: efficient multicellular movement. Prior to protostome-deuterostome divergence, the bilaterian animal likely acquired sensorimotor specializations, including divisions between cardiac, skeletal/somatic and visceral muscles, which are today regulated by similar gene families in Drosophila and vertebrates e.g. *mef2* genes (Taylor and Hughes, 2017). Skeletal myogenesis is initiated during gastrulation, shortly after mesendoderm formation, but conserved bilaterian pathways leading specifically to skeletal muscle (as opposed to cardiac or visceral) have been harder to discern. One likely reason for this is that new regulatory steps have evolved in each lineage since their divergence.

A key step in vertebrate evolution was the chordate transition through which animals acquired the notochord, post-anal tail, gill slits and a more complex dorsal nerve cord, facilitating swimming (Brunet et al., 2015; Gee, 2018; Gerhart, 2001; Satoh et al., 2012). Throughout vertebrates, the notochord patterns the neural tube and paraxial mesodermal tissue by secreting Hedgehog (Hh) signals that promote motoneuron and early muscle formation (Beattie et al., 1997; Blagden et al., 1997; Du et al., 1997; Münsterberg et al., 1995; Roelink et al., 1994). Nevertheless, in the absence of either notochord or Hedgehog signalling, muscle is formed in vertebrate somites (Blagden et al., 1997; Dietrich et al., 1999; Du et al., 1997; Grimaldi et al., 2004; Zhang et al., 2001). How might deuterostome muscle have formed prior to evolution of the notochord?

A change in function of the *Tbxt* gene, a T-box (Tbx) family paralogue, occurred during chordate evolution such that *Tbxt* now directly controls formation of posterior mesoderm, notochord and post-anal tail in vertebrates (Chiba et al., 2009; Showell et al., 2004). Hitherto, *Tbxt* may have distinguished ectoderm from endoderm and regulated formation of the most posterior mesendoderm (Arenas-Mena, 2013; Kispert et al., 1994; Woollard and Hodgkin, 2000; Yasuoka et al., 2016). In vertebrates, *Tbxt* genes also promote slow myogenesis (Coutelle et al., 2001; Halpern et al., 1993; Martin and Kimelman, 2008; Weinberg et al., 1996). Other Tbx genes, such as *Tbx1*, *Tbx4*, *Tbx5*, *Tbx16* and *Tbx6*, also influence sarcomeric muscle development (Chapman and Papaioannou, 1998; Griffin et al., 1998; Hasson et al., 2010; Kimmel et al., 1989; Manning and Kimelman, 2015; Weinberg et al., 1996; Windner et al., 2015). For example, the Tbx6 family is implicated in early stages of paraxial mesoderm commitment, somite patterning and the generation and positioning of muscle precursor cells (Bouldin et al., 2015; Chapman and Papaioannou, 1998; Kimmel et al., 1989; Manning and Kimelman, 2015; Nikaido et al., 2002; White et al., 2003; Windner et al., 2012). It is unclear, however, whether the Tbx genes promote myogenesis directly, and/or are required for earlier events in mesoderm development that are necessary for subsequent myogenesis.

In vertebrates, a key essential step in skeletal myogenesis is activation of myogenic regulatory factors (MRFs) encoded by the *myf5* and *myod* genes (Hinits et al., 2009; Hinits et al., 2011; Rudnicki et al., 1993). Distinct myogenic cell populations initiate *myf5* and *myod* expression in different ways, the genes being induced by distinct signals through distinct *cis*-regulatory elements in different skeletal muscle precursor cells (Buckingham and Rigby, 2014). As the anteroposterior axis forms and extends, *de novo* induction of *myf5* and *myod* mRNAs in slow and fast muscle precursors occurs in tissue destined to generate each successive somite. Once expressed, these MRF proteins have two functions: to remodel chromatin and directly enhance transcription of muscle genes (reviewed in (Buckingham and Rigby, 2014). In zebrafish, myogenesis begins at about 75% epiboly stage when adaxial cells that flank the shield/organizer/nascent notochord (hereafter called pre-adaxial cells; diagrammed in Fig. 1A), begin MRF expression (Hinits et al., 2009; Melby et al., 1996). Pre-adaxial cells express both *myf5* and *myod*, converge to form two rows of adaxial cells flanking the notochord, become incorporated into somites and differentiate into slow muscle fibres (Coutelle et al., 2001; Devoto et al., 1996; Weinberg et al., 1996). Loss of either Myf5 or Myod alone is not sufficient to prevent slow myogenesis, but loss of both completely inhibits adaxial slow muscle formation (Hinits et al., 2009; Hinits et al., 2011). Dorsal tissue immediately lateral to the pre-adaxial cells, the paraxial mesoderm (Fig. 1A), expresses *myf5* but little *myod* and subsequently generates fast muscle once somites have formed, upregulating *myod* in the process. A key to understanding myogenesis in both paraxial and adaxial cells is thus the mechanism(s) by which *myf5* and *myod* expression is regulated.

**Fig. 1.**
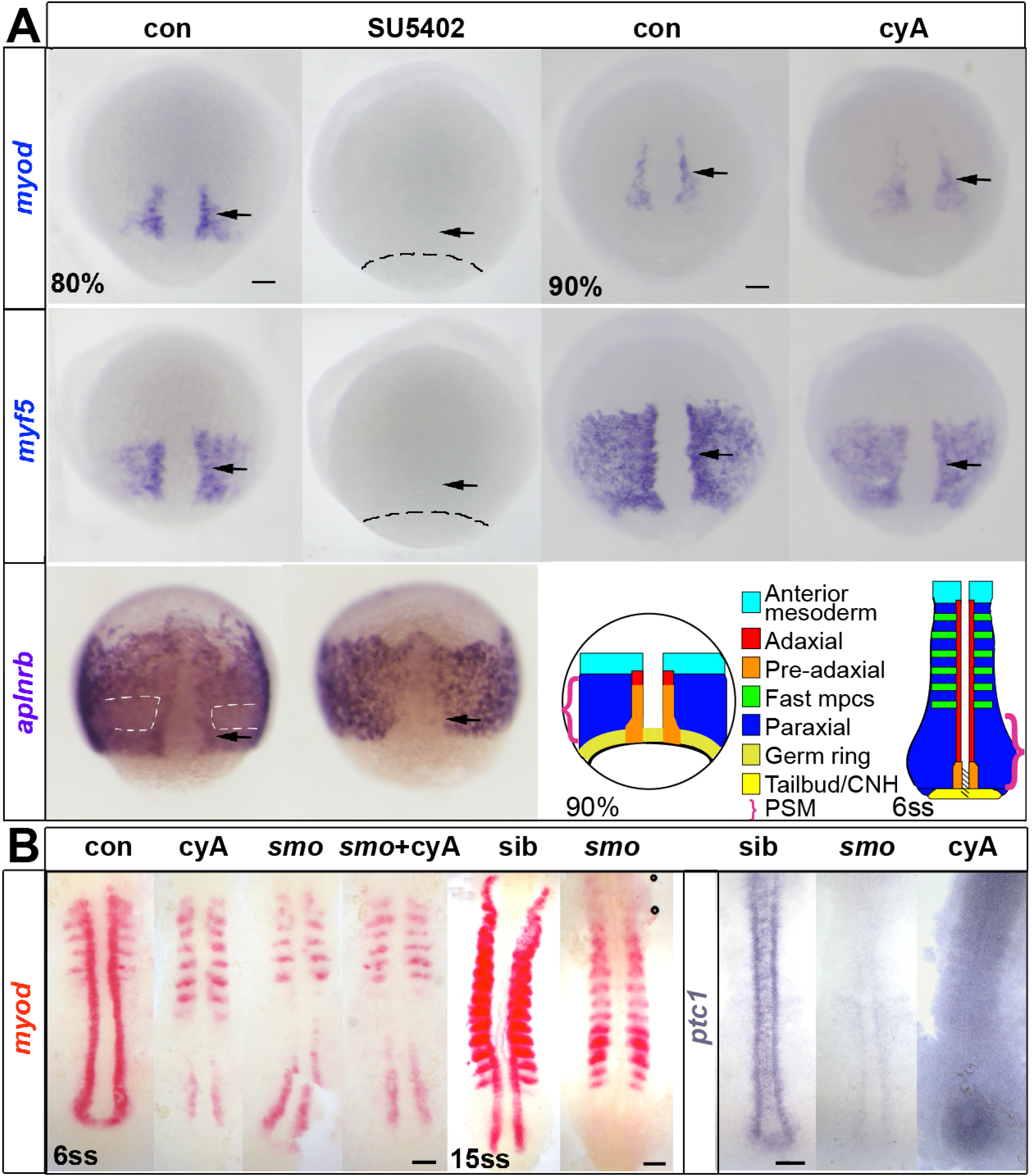
Inhibition of Fgf signalling blocks initial induction of *myod* and *myf5* expression. *In situ* mRNA hybridization for *myod* and *myf5* in control untreated, cyclopamine-treated (cyA, 100 μM), SU5402-treated and double SU5402- and cyA-treated wild type or mutant embryos, shown in dorsal view, anterior to top. **A.** Adaxial (arrows) and paraxial *myod* and *myf5* mRNAs are lost upon SU5402-treatment (60 µM) from 30% to 80 or 90% epiboly (dashes indicate approximate position of germ ring) but are unaffected by cyA treatment. The anterior mesoderm marker *aplnrb* is normally down-regulated in paraxial presomitic cells expressing *myf5* (white dashes) and up-regulated in adaxial cells (arrow). Both changes were absent after SU5402-treatment. Schematics illustrate the location of equivalent cell types at two successive stages. mpcs = muscle precursor cells, CNH = chordoneural hinge (hatched), PSM = presomitic mesoderm (brackets). **B.** *Smo^b641^* mutants retain pre-adaxial *myod* mRNA at 6ss even after cyA treatment, but lack pre-adaxial *myod* mRNA at 15ss. *Ptc1* mRNA downregulation shows that both *smo* mutation and cyA treatment (shown after longer colour reaction) fully block Hh signalling throughout the axis. Bars: 50 µm.

Intrinsic factors such as Tbx proteins likely interact with extrinsic positional cues within the embryo to pattern myogenesis. Fgf and Tbx function have long been known to interact to drive early mesendoderm patterning, but how directly they control early embryonic myogenesis remains unclear (Kimelman and Kirschner, 1987; Showell et al., 2004; Slack et al., 1987). Various Fgf family members are expressed close to myogenic zones during vertebrate gastrulation (Isaacs et al., 2007; Itoh and Konishi, 2007; Wilkinson et al., 1988). In zebrafish, Fgf signalling is required for mesendoderm formation, tailbud outgrowth and normal fast myogenesis (Draper et al., 2003; Griffin et al., 1995; Groves et al., 2005; Reifers et al., 1998; Yin et al., 2018). Fgf signalling is also thought to be involved in early expression of *myf5* and *myod* in pre-adaxial cells, but the mechanism of initial induction of *myf5* and *myod* is unknown (Ochi et al., 2008). Expression of *fgf3*, *fgf4*, *fgf6a*, *fgf8a* and *fgf8b* has been detected in the chordoneural hinge (CNH, Fig. 1A) adjacent to pre-adaxial cells (Draper et al., 2003; Groves et al., 2005; Thisse and Thisse, 2005). Subsequently, Hedgehog (Hh) signalling from the ventral midline maintains MRF expression and progression of the pre-adaxial cells into terminal slow muscle differentiation (Coutelle et al., 2001; Hirsinger et al., 2004; Lewis et al., 1999). Here, we show how both Fgf and Hh extracellular signals cooperate with Tbx genes to control fast and slow myogenesis. In the trunk, Fgf signalling is required for the initiation of myogenesis and acts in cooperation with Tbx16/Tbxta function directly on *myf5* and *myod*. In the tail, by contrast, direct MRF gene induction by Fgf is not required and the evolutionary novelty of midline-derived Hh-signalling accounts for adaxial myogenesis.

## Results

### Fgf signalling is essential for induction of adaxial myf5 and myod expression

Adaxial myogenesis is driven by successive Fgf and Hh signals. When Hh signalling was prevented with the Smo antagonist cyclopamine (cyA), *myf5* and *myod* mRNAs in pre-adaxial cells were unaffected at 90% epiboly (Fig. 1A). In contrast, when Fgf signalling was inhibited with SU5402 both *myf5* and *myod* mRNAs were lost (Fig. 1A)(Ochi et al., 2008). To show that lack of MRFs was not due to failure of gastrulation caused by SU5402-treatment, we analysed expression of *aplnrb* mRNA, an anterior mesodermal marker (Zeng et al., 2007). At 80% epiboly, *aplnrb* mRNA has a complex and informative expression pattern, marking the anterior invaginating mesoderm cells around the germ ring and the pre-adaxial cells, but appears down-regulated in more lateral regions expressing *myf5* but not *myod* (Figs 1A; S1, see Table S1 for quantification of this and subsequent experiments). In SU5402-treated embryos, *aplnrb* mRNA reveals the normal invagination of mesoderm and *aplnrb*-expressing cells flanking the organiser. Both the downregulation of *aplnrb* mRNA in paraxial trunk mesoderm that normally expresses *myf5* alone, and pre-adaxial *aplnrb* up-regulation were absent in SU5402-treated embryos (Fig. 1A). Thus, early Fgf signalling is required for the initiation of skeletal myogenesis in future trunk regions.

As trunk myogenesis proceeds, Hh signalling becomes essential for adaxial slow myogenesis. At 6ss, in *smo* mutants, cyA-treated embryos and even cyA-treated *smo* mutants, all of which lack Hh signalling, *myod* mRNA is lost from adaxial slow muscle but persists in paraxial fast muscle precursors (Fig. 1B). Nevertheless, *myod* mRNA transiently accumulates in pre-adaxial cells of presomitic mesoderm (PSM) destined to make trunk somites, but is then lost in anterior PSM (Fig. 1B)(Barresi et al., 2000; Lewis et al., 1999; Osborn et al., 2011; van Eeden et al., 1996; van Eeden et al., 1998). Thus, as suggested previously (Coutelle et al., 2001; Ochi et al., 2008), in the wild type situation trunk pre-adaxial *myod* expression is maintained and enhanced by Hh. In contrast, during tail myogenesis at 15ss and thereafter, no pre-adaxial *myod* expression was detected in *smo* mutants or cyA-treated embryos (Fig. 1B and data not shown). These data suggest that whereas Hh is necessary for induction of adaxial myogenesis in the tail, Fgf-like signals initiate *myod* expression in trunk pre-adaxial cells.

Additional evidence emphasizes the greater importance of Hh in tail myogenesis. In *shha* mutants, slow muscle is lost from tail but remains present in trunk somites (Fig. 2A). Moreover, injection of *myod* or *myog* mRNA into embryos lacking Hh signalling can rescue slow myogenesis in trunk but not in tail (Fig. S2). Similarly, absence of notochord-derived signals in *noto* (*flh*) mutants, in which the nascent notochord loses notochord character and converts to muscle (Coutelle et al., 2001; Halpern et al., 1995), ablates tail but not trunk slow muscle (Fig. 2B). Treatment of *noto* mutants with cyA shows that *myod* expression is initiated in trunk pre-adaxial cells adjacent to the transient pre-notochordal tissue, but fails to be maintained owing to the blockade of floorplate-derived Hh signals (Fig. 2C). Taken together, these data show that Hh can initiate and then maintain MRF gene expression, but that other signals initiate slow myogenesis in the trunk.

**Fig. 2.**
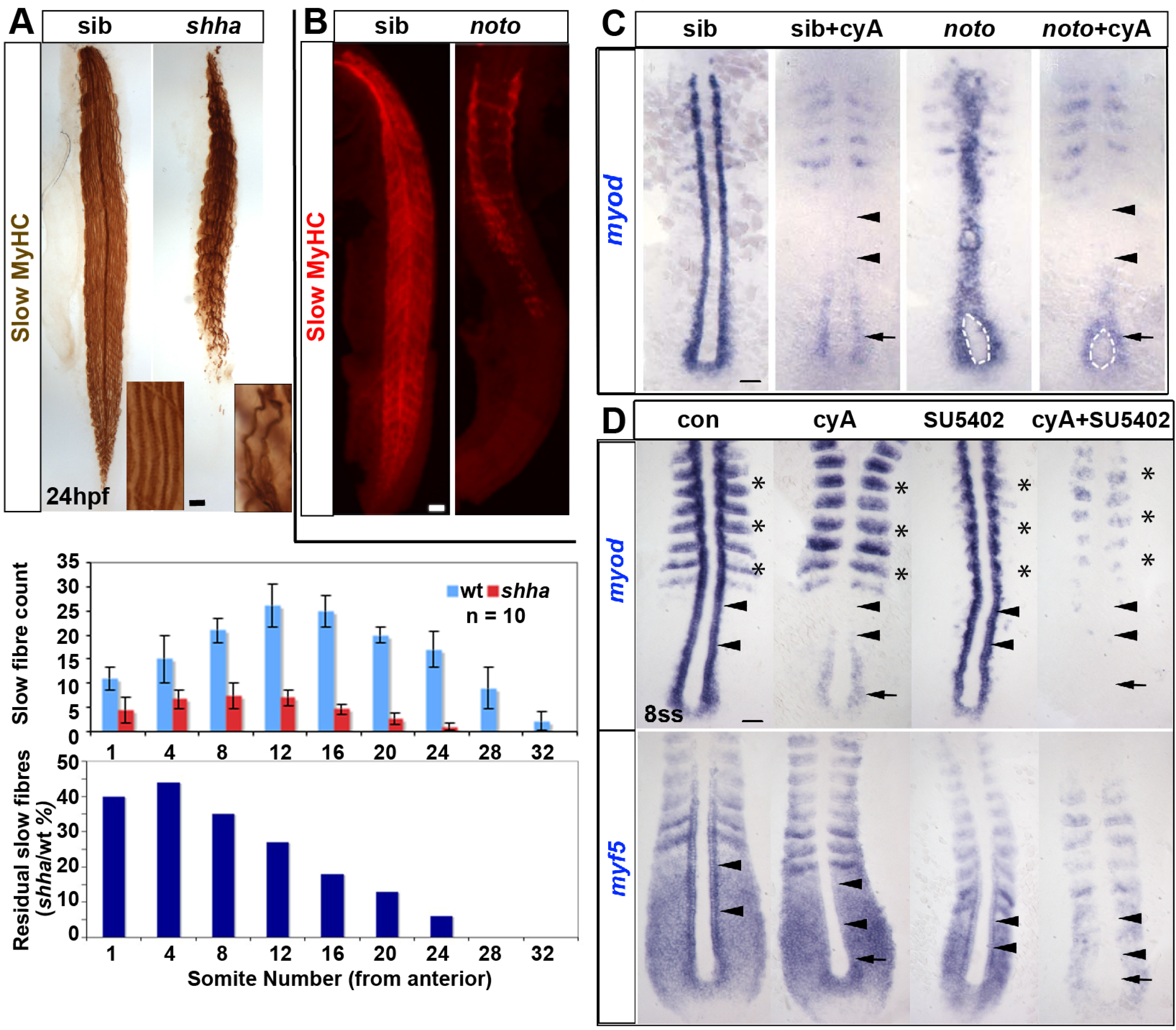
Successive Fgf and Hh signals drive trunk slow myogenesis. Immunodetection of slow MyHC in *shha* and *noto* mutants. **A.** Reduction in slow fibres in *shha* mutants is greater in tail than in trunk. Insets show individual fibres magnified. Upper graph shows mean ± SD of slow fibre number in the indicated somites from ten independent embryos of each genotype. Lower graph shows fraction of wild type fibres remaining in mutant. **B.** Trunk-specific residual slow muscle in *noto* mutant. **C.** 5-somite stage (5ss) embryos from a *noto^n1^* heterozygote incross treated with cyA at 30% epiboly stage, showing loss of adaxial *myod* mRNA in anterior presomitic mesoderm (arrowheads), but retention in the most posterior pre-adaxial mesoderm (arrows) flanking the chordoneural hinge (white outline). **D.** SU5402 (50 µM) from tailbud stage ablates residual pre-adaxial *myod* and *myf5* mRNAs in cyA-treated 8ss embryos (arrows). Expression of paraxial *myod* in fast muscle precursors (asterisks) is not affected by cyA but is decreased by SU5402. Bars: 50 µm.

We next tested whether Hh-independent *myod* expression and *myf5* up-regulation in trunk pre-adaxial cells requires Fgf signalling. Treatment with cyA left residual pre-adaxial *myod* and *myf5* mRNA flanking the base of the notochord at trunk levels, but ablated adaxial expression in slow muscle precursor cells (Fig.2D). The residual expression was ablated when, in addition to cyA, SU5402 was used to block *fgf* signalling from 30% epiboly (Fig. 2D). Application of SU5402 alone diminished *myf5* and *myod* mRNA accumulation up to tailbud stage, but caused little if any reduction of adaxial *myf5* and *myod* mRNAs in the tailbud region at the 6 somite stage (6ss), after midline *shha* function had commenced (Fig. 1B,C)(Krauss et al., 1993). Nevertheless, SU5402 greatly diminished *myod* expression in somitic fast muscle precursors and reduced the extent of *myf5* expression in paraxial PSM (Fig. 1B,C)(Groves et al., 2005; Reifers et al., 1998).

### Fgf3, fgf4, fgf6a and fgf8a collaborate to promote MRF expression

To identify candidate Fgf regulators of pre-adaxial myogenesis, the expression patterns of *fgf3*, *fgf4*, *fgf6a* and *fgf8a* were investigated on wild type embryos (Fig. S3A). As reported previously, *fgf4*, *fgf6a* and *fgf8a* mRNAs were all detected in the posterior dorsal midline at 80% epiboly, followed by *fgf3* in the chordoneural hinge (CNH) and posterior notochord (Fig. S3A)(Kudoh et al., 2001; Thisse and Thisse, 2005; Yamauchi et al., 2009). The temporal and spatial expression of *fgf3*, *fgf4*, *fgf6a* and *fgf8a* during gastrulation and early somitogenesis make them excellent candidates for Fgf regulators of *myf5* and *myod*.

To evaluate the role of Fgfs in *myf5* and *myod* expression, each Fgf was knocked down with previously validated antisense morpholino oligonucleotides (MOs) in wild type embryos (Fig. S3B). At 80% epiboly, there was little or no decrease of *myf5* or *myod* mRNA in individual *fgf* morphants or *fgf8a* mutant embryos (Fig. S3B). Combinatorial knockdown of several fgfs led to progressively more severe loss of *myod* mRNA and reduction of the raised pre-adaxial and paraxial levels of *myf5* mRNA (Fig. 3A). Thus, specific Fgfs collaborate to drive the initial expression of *myod* and *myf5* in pre-adaxial and paraxial cells.

**Fig. 3.**
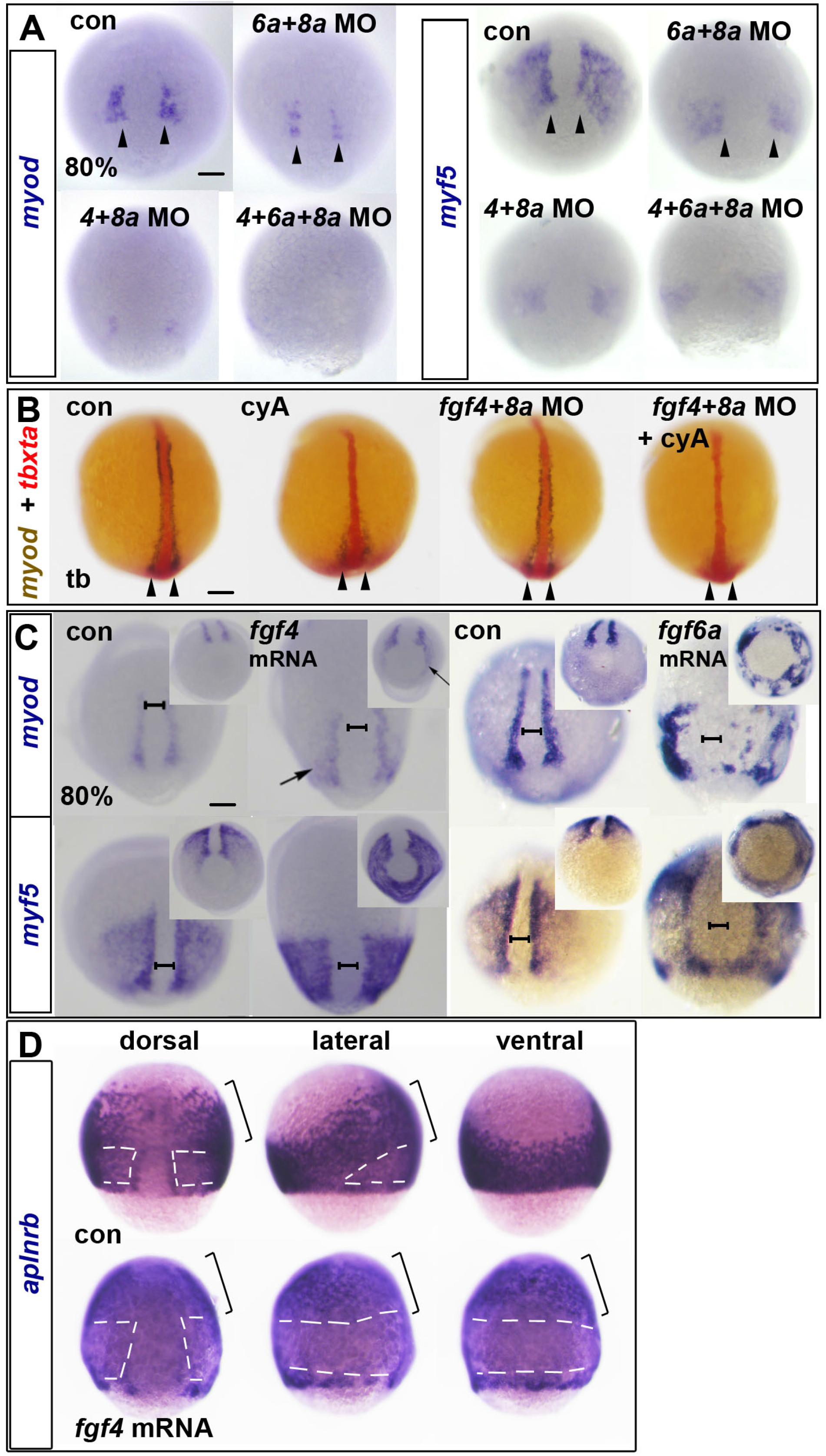
Dorsally-expressed Fgfs drive paraxial myogenesis. In situ mRNA hybridization for *myod* and *myf5* (A,C) or *aplnrb* (D) mRNAs at 80% epiboly or *tbxta* (red) and *myod* (blue/brown) at tailbud stage (tb) (B). **A.** Reduction of *myod* and *myf5* mRNAs in *dual and triple fgf* MO-injected wild type embryos. Arrowheads indicate nascent adaxial cells. B. In contrast to 80% epiboly (compare Fig. S3C), at tailbud stage, cyclopamine (cyA) treatment ablates anterior adaxial *myod* mRNA, but leaves pre-adaxial expression intact (arrowheads). Injection of *fgf4*+*fgf8a* MOs ablate residual *myod* mRNA. **C.** Fgf4 or Fgf6a mRNA injection up-regulates *myod* and *myf5* mRNAs around the marginal zone. Note the widening of the unlabelled dorsal midline region (brackets). Insets show the same embryos viewed from vegetal pole. **D.** Fgf4 mRNA injection down-regulates *aplnrb* mRNA around the marginal zone (white dashes) but not anteriorly (brackets) in the dorsalized embryo. Bars: 100 µm.

By tailbud stage, however, *fgf4*+*fgf8a* MO treatment alone had little effect on *myod* mRNA accumulation, presumably due to the presence of Hh in the midline (Fig. 3B). Congruently, cyA-treatment reduced anterior adaxial *myod* mRNA, but pre-adaxial expression persisted after blockade of Hh signalling (Fig. 3B). Pre-adaxial *myod* mRNA was ablated by cyA-treatment of embryos injected with *fgf4+fgf8a* MO (Fig. 3B). Control injection of *fgf4*+*fgf8a* MOs into vehicle-treated embryos did not reduce *myod* mRNA accumulation, confirming that Hh can initiate trunk adaxial *myod* expression (Fig. 3B). Thus, expression of Fgf4 and Fgf8a in the shield, CNH and posterior notochord, provides a spatiotemporal cue for pre-adaxial myogenic initiation in the tailbud.

To test the ability of Fgfs to promote myogenesis further, we generated ectopic Fgf signals by injection of *fgf4* or *fgf6a* mRNA into wild type embryos, and analysed *myf5* and *myod* mRNA at 80% epiboly (Fig. 3C). Both *myod* and *myf5* mRNAs were upregulated in more ventral regions at levels comparable to those in adaxial cells of control embryos, despite the absence of Hh signalling in these regions. Over-expression of *fgf4* mRNA upregulated *myf5* mRNA in an initially uniform band around the embryo that extended towards the animal pole for a distance similar to that of *myod* mRNA in adaxial cells of controls (Fig. 3C). *Myod* was less easily induced, but in similar regions of the mesoderm. The posterior notochord and shield still lacked MRFs and appeared wider. However, this dorsalmost tissue was not enlarged; the number of notochord cells scored by DAPI stain appeared normal (data not shown). The embryos became ovoid, with a constriction around the germ ring that appeared to stretch and broaden the notochord. There was a positive correlation between the extent of *myf5* and *myod* mRNA up-regulation and the extent of deformation towards egg-shape. *Aplnrb* mRNA persisted in animal regions of the mesoderm, but was downregulated where *myf5* mRNA was induced nearer to the margin, revealing that anterior/cranial mesoderm is present but resistant to Fgf-driven MRF induction and *aplnrb* suppression (Fig. 3D). Thus, Fgf4 dorsalized the embryo, converting the entire posterior paraxial and ventral mesoderm to a myogenic profile with some regions expressing only *myf5* and others expressing also *myod*, particularly around the germ ring. Fgf6a overexpression also induced ectopic MRFs in cells around the germ ring, which then appeared to cluster (Fig. 3C). Taken together, these data show that posterior/dorsal Fgf signals initiate MRF expression in both pre-adaxial slow and paraxial fast muscle precursors in pre-somitic mesoderm.

### MRFs are initially induced by fgfs, tbxta and tbx16

Zebrafish Tbx genes, including *tbxta* and *tbx16* (formerly called *no tail* (*ntla*) and *spadetail* (*spt*), respectively), are potentially important mediators of Fgf signalling in gastrulating embryos (Amaya et al., 1993; Griffin et al., 1995; Smith et al., 1991; Sun et al., 1999). *Tbx16* is suppressed by a dominant negative Fgf receptor (FgfR) (Griffin et al., 1998). However, whether *tbxta* and *tbx16* activities are altered by SU5402 treatment, which also blocks FgfR function, is unclear (Rentzsch et al., 2004; Rhinn et al., 2005). Wild type embryos at 30% epiboly were therefore exposed to SU5402 and subsequently fixed at 80% epiboly or 6ss to investigate expression of *tbxta and tbx16* (Fig. 4A). Compared to controls, embryos treated with 10 μM SU5402 showed diminished *tbxta* expression in notochord and less *tbxta* and *tbx16* in the germ ring, particularly in dorsal paraxial regions. At 6ss, expression of *tbxta* and *tbx16* was mildly reduced in tailbud by 10 μM SU5402 (Fig. 4A). When the concentration of SU5402 was increased to 30 μM, expression of *tbxta* and *tbx16* was abolished throughout the trunk (Fig. 4A). Thus, Fgf-like signalling is required for normal Tbx gene expression in the mesoderm.

**Fig. 4.**
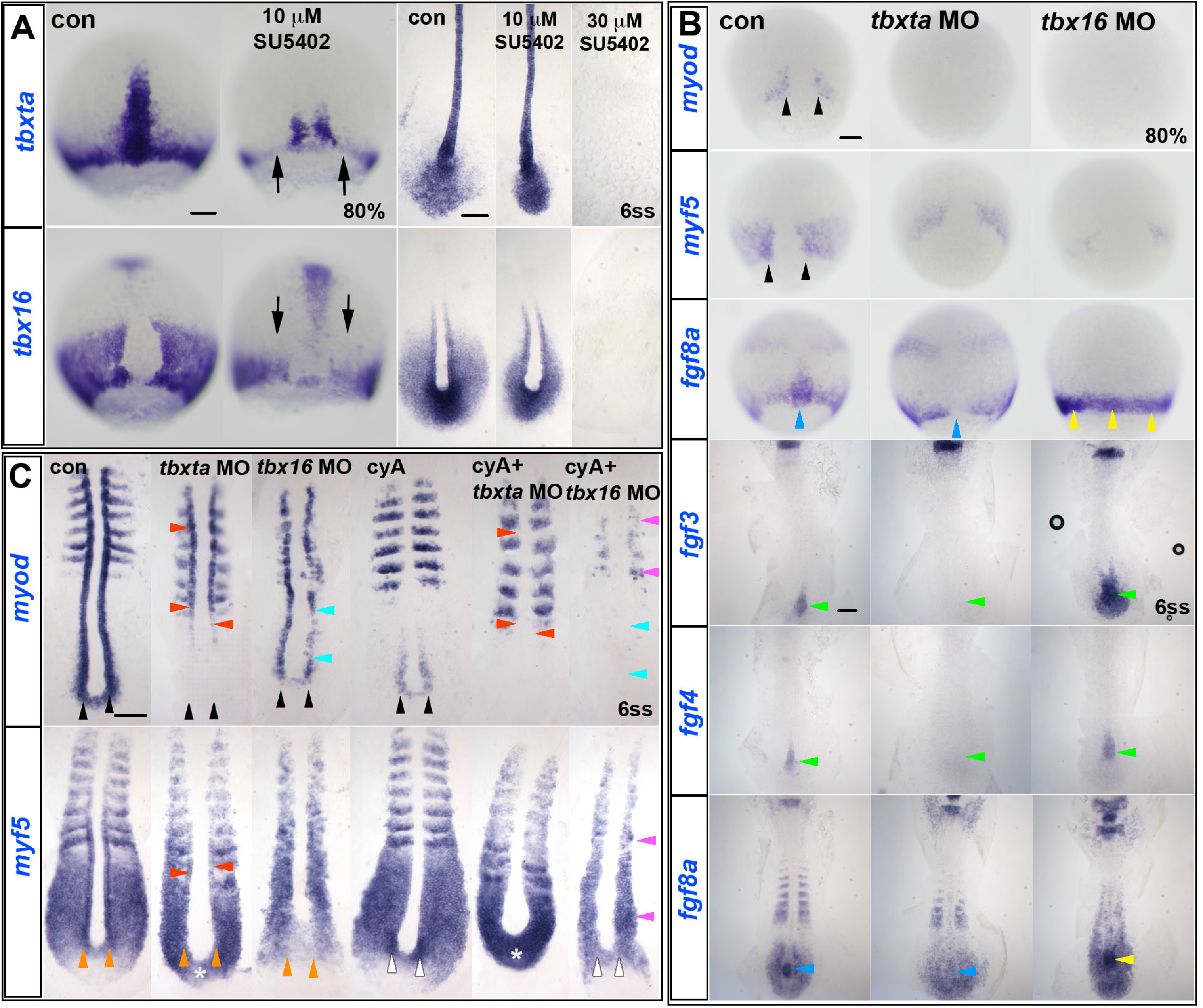
Redundant Fgf/Tbx and Hh signals required for MRF induction. In situ mRNA hybridization for *tbxta* and *tbx16* in control untreated and SU5402-treated from 30% embryos (A), for *myod, myf5, fgf8a*, *fgf*3 and *fgf4*, in control, *tbxta* MO-and *tbx16* MO-injected wild type (wt) embryos (B), and for *myod* and *myf5* in control, *tbxta* MO-and *tbx16* MO-injected wt embryos treated with or without 0.1 mM cyA (C). **A.** In 10 µM SU5402-treated wt embryos, *tbxta* and *tbx16* transcripts are decreased (arrows) at 80% epiboly, but almost normal at 6ss. Both transcripts are absent in 30 µM SU5402-treated embryos at 6ss. **B.** Adaxial *myod* expression (black arrowheads) is completely ablated in *tbxta* or *tbx16* morphants at 80% epiboly, and *myf5* expression is greatly decreased. *Fgf8a* mRNA is ablated in posterior notochord of *tbxta* morphants (blue arrowheads), but upregulated in *tbx16* morphants around the germ marginal zone at 80% and in posterior notochord at 6ss (yellow arrowheads). Expression of *fgf3* and *fgf4* is absent in posterior notochord of *tbxta* morphants, but enhanced in that location in *tbx16* morphants (green arrowheads). **C.** At 6ss, pre-adaxial *myod* expression (black arrowheads) is lost in *tbxta* morphant tailbud, and diminished in *tbx16* morphants. Injection of *tbx16* MO, but not *tbxta* MO, reduces adaxial *myf5* mRNA to the level in paraxial mesoderm (orange arrowheads), whereas *tbxta* MO but not *tbx16* MO up-regulates *myf5* mRNA in posterior tailbud (asterisks). *Tbx16* MO abolishes pre-adaxial *myf5* mRNA in cyA-treated embryos (white arrowheads). Adaxial *myf5* and *myod* transcripts recover in *tbxta* morphants, but are ablated by cyA-treatment (red arrowheads). CyA-treatment of *tbx16* morphants ablates adaxial *myod* expression throughout the axis (cyan arrowheads), leaving only residual paraxial *myod* and *myf5* expression (mauve arrowheads). Bars: 100 µm.

Tbxta is required for normal *myf5* and *myod* expression in posterior regions during tailbud outgrowth, partly due to loss of Hh signalling from the missing notochord (Coutelle et al., 2001; Weinberg et al., 1996). To test whether Tbx genes are required for MRF expression at earlier stages, each Tbx gene was knocked down and MRF and Fgf expression analysed at 80% epiboly. Tbxta knockdown reduced expression of dorsal midline Fgfs, ablated *myod* mRNA and reduced *myf5* mRNA accumulation in pre-adaxial cells to the level in paraxial regions (Fig. 4B). However, Tbxta knockdown had little effect on either *fgf8a* or *myf5* mRNAs in more lateral paraxial mesoderm or the germ ring (Fig. 4B). This correlation raised the possibility (addressed below) that Tbxta may drive MRF expression through induction of Fgf expression. Tbx16 knockdown, on the other hand, ablated pre-adaxial *myod* mRNA and reduced both pre-adaxial and paraxial *myf5* mRNA without reduction of Fgf expression (Fig. 4B). Indeed, both germ ring *fgf8a* mRNA at 80% epiboly and dorsal midline *fgf3*, *fgf4* and *fgf8a* mRNAs in the tailbud at 6ss appeared increased (Fig. 4B), as previously described (Warga et al., 2013). As Tbx16 expression persists in *tbxta* mutants (Amack et al., 2007; Griffin et al., 1998), these data raise the possibility that Tbx16 is required to mediate the action of Fgf signals on myogenesis.

*Tbx16* null mutants show a failure of convergent migration of mesodermal cells into the paraxial region (Ho and Kane, 1990; Molven et al., 1990) which, by reducing mesodermal cells flanking the CNH, may contribute to the reduction in MRF mRNAs observed at 80% epiboly. Nevertheless, *tbx16* mutants generate enough paraxial mesoderm that reduced numbers of both paraxial fast muscle and adaxially-derived slow muscle fibres arise after Hh signalling commences (Honjo and Eisen, 2005; Weinberg et al., 1996). To investigate whether Tbx16 is required for initial induction of *myf5* and/or *myod* expression in pre-adaxial cells, we titrated MO to reduce *tbx16* function to a level that did not prevent accumulation of significant numbers of trunk mesodermal cells and examined *myf5* and *myod* expression at 6ss (Fig. 4C). In *tbx16* morphants, *myod* mRNA was readily detected in adaxial cells adjacent to notochordal Hh expression (Fig. 4C, cyan arrowheads). Treatment of these *tbx16* morphants with cyA to block Hh signalling, however, completely ablated adaxial *myod* expression, leaving only weak *myod* in paraxial somitic fast muscle precursors (Fig. 4C, cyan and purple arrowheads). In contrast, treatment of control embryos with cyA left pre-adaxial *myod* mRNA intact (Fig. 4C, black arrowheads). These findings show that Fgf-driven pre-adaxial *myod* expression flanking the CNH requires Tbx16 function.

Adaxial *myf5* expression also requires Tbx16. Tbx16 knockdown reduces *myf5* mRNA accumulation in the posterior tailbud, and also diminishes the upregulation of *myf5* mRNA in pre-adaxial and adaxial cells (Fig. 4C, orange arrowheads). Addition of cyA to *tbx16* morphants has little further effect on *myf5* expression (Fig. 4C, white arrowheads). In contrast, cyA treatment alone reduces adaxial *myf5* mRNA in anterior PSM, but does not affect the *myf5* up-regulation in pre-adaxial cells or tailbud *myf5* expression (Fig. 4C, white arrowheads). Additional knockdown of Tbx16 in cyA-treated embryos prevents pre-adaxial *myf5* up-regulation (Fig. 4C, white arrowheads). Thus, Tbx16 is required for Fgf to up-regulate both *myf5* and *myod* in pre-adaxial cells.

Both pre-adaxial and anterior PSM adaxial *myod* expression were also absent in *tbxta* morphants, but recovered in somites, again due to midline-derived Hh signalling (Fig. 4C and (Coutelle et al., 2001)). In marked contrast, Tbxta knockdown up-regulated *myf5* mRNA in tailbud (Fig. 4C, asterisk), presumably reflecting loss of tailbud stem cells that lack *myf5* mRNA. Taken together, the data strongly suggest that *tbx16* is required for midline-derived Fgf signals to induce *myod* and up-regulate *myf5* in pre-adaxial cells in tailbud. In contrast, the loss of MRF expression in *tbxta* mutants could be simply explained by loss of midline-derived Fgf signals, and/or might require some other Tbxta-dependent process.

### Myf5 and myod induction by Fgf signalling require Tbx16

To test rigorously the idea that Tbx16 is required for Fgf to induce MRFs, *fgf4* mRNA was injected into embryos from a *tbx16* heterozygote incross. Whereas Fgf4 up-regulated *myf5* and *myod* mRNAs all around the germ ring in siblings, in sequence-genotyped *tbx16* mutant embryos no up-regulation was detected (Fig. 5A). It is clear that mesoderm was present in *tbx16* mutants because the mRNAs encoding Aplnrb, Tbxta, Tbx16 and Tbx16-like (formerly Tbx6-like) are present in *tbx16* mutants (Fig. S4; (Griffin et al., 1998; Morrow et al., 2017)). The effect of Fgf4 does not act by radically altering *tbxta* or *tbx16* gene expression (Fig. 5B). Thus, Tbx16 is required for Fgf-driven expression of MRFs in pre-somitic mesoderm.

**Fig. 5.**
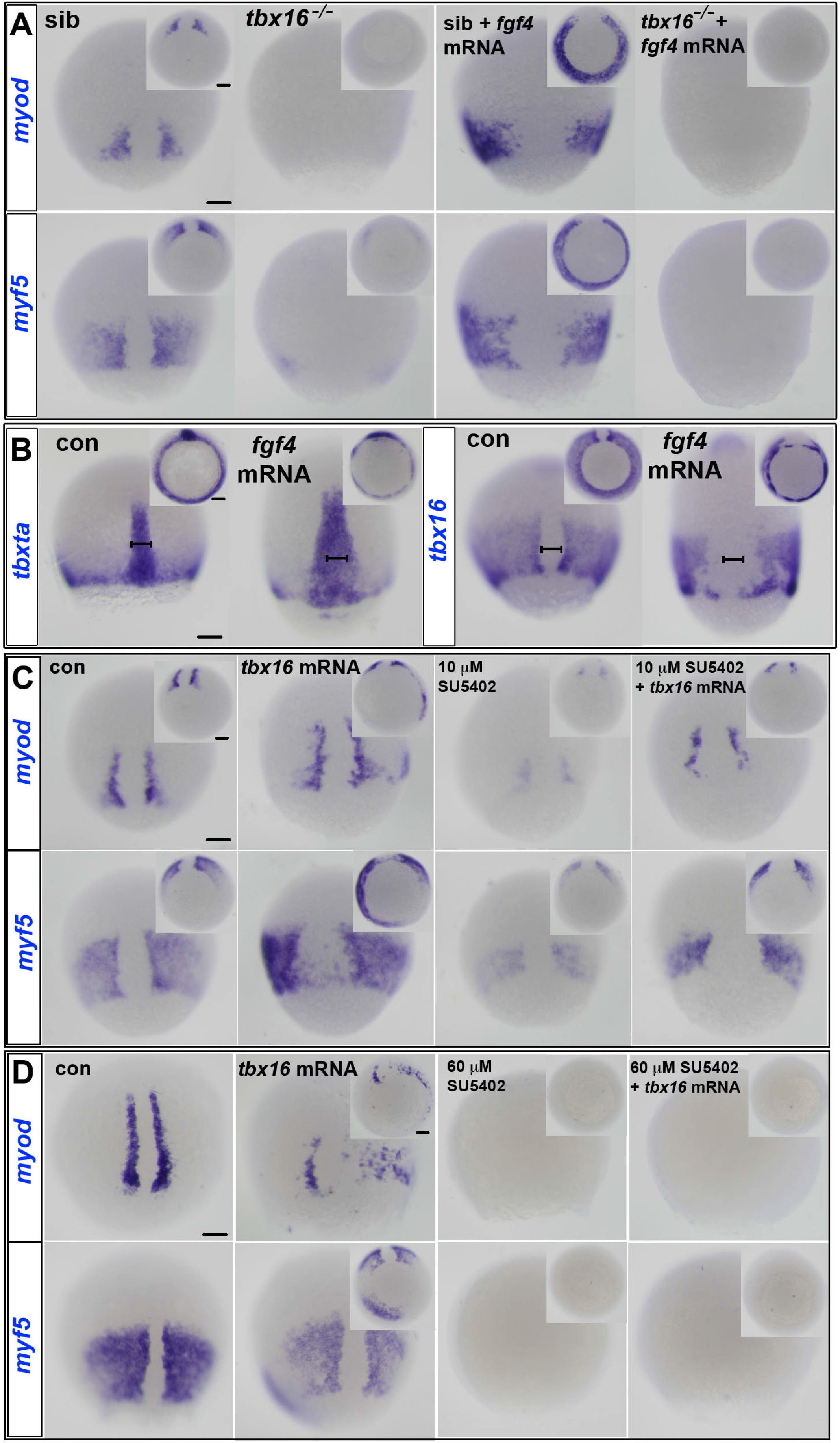
Tbx16 is necessary and sufficient for MRF induction. *In situ* mRNA hybridization for the indicated mRNAs of control uninjected and *fgf4* (A,B) or *tbx16* (C,D) mRNA-injected embryos at 80% epiboly stage in *tbx16^+/-^* incross (A) and wild type (B-D). Dorsal views. Insets ventral views. **A.** *Myf5* and *myod* mRNAs flank the dorsal midline in siblings (sib), but are absent or greatly diminished in *tbx16^-/-^* mutants. *Fgf4* widened notochord (bars) and induced ectopic *myod* and *myf5* mRNA around the germ marginal zone of siblings, but did not rescue expression in *tbx16^-/-^* embryos. **B.** *Tbxta* mRNA reveals widened notochord (bars) in wild type embryos injected with *fgf4* mRNA. Both *tbxta* and *tbx16* mRNAs show clumping in the germ ring after overexpression of *fgf4*. **C,D.** *In situ* mRNA hybridization at 80% epiboly stage for *myf5* and *myod* mRNAs in wild type control or *tbx16* mRNA-injected embryos treated with SU5402 at 10 µM (C) or 60 µM (D). *Myf5* and *myod* mRNAs are ectopically induced in posterior mesoderm by Tbx16 expression, decreased by administration of 10 µM SU5402 in wild type, and rescued in SU5402-treated embryos by overexpression of *tbx16*. High dose SU5402 prevents MRF expression, even after *tbx16* mRNA injection. Bars: 100 µm.

### Tbx16 requires Fgf-like signalling to rescue myf5 and myod expression

The results so far show that *tbx16* function is necessary for Fgf to induce *myf5* and *myod* (Fig. 5A). To determine if increased Tbx16 activity is sufficient to induce MRFs, Tbx16 was overexpressed. Injection of *tbx16* mRNA caused ectopic expression of both *myf5* and *myod* in the germ marginal zone (Fig. 5C). Notably, Tbx16 over-expression induced *myf5* mRNA in a much broader region than *myod* mRNA.

We next wanted to determine whether Tbx16 could induce expression of *myf5* or *myod* in the absence of Fgf signalling. Exposure to high dose SU5402, which down-regulates endogenous *tbx16* and *tbxta* mRNA (Fig. 4A), prevented MRF induction by injection of *tbx16* mRNA (Fig. 5D). Nevertheless, when *tbx16* mRNA was injected into low dose (10 μM) SU5402-treated embryos, which normally have reduced MRF expression, the level of *myf5* and *myod* mRNAs was rescued (Fig. 5C). However, *tbx16* mRNA was less effective at ectopic MRF induction in the presence of SU5402 (Fig. 5C). These results demonstrate that Fgf signalling cooperates with Tbx16 activity in inducing expression of *myf5* and *myod* in pre-adaxial cells at gastrulation stages. Moreover, Tbx16 requires Fgf-like signals to induce MRF gene expression.

### Myf5 and Myod are direct transcriptional targets of Tbx16

In order to determine further the regulatory relationship between Tbx16 and *myf5* and *myod* we interrogated ChIP-seq experiments for endogenous Tbx16 on 75-85% epiboly stage embryos (see Materials and Methods)(Nelson et al., 2017). Our analyses revealed a highly significant peak at −80 kb upstream (*myf5* Distal Element, 5DE) and two binding peaks proximal to *myf5* (Proximal Elements, 5PE1,5PE3) (Fig. 6A,B and Supplementary Table S3). To determine which of these binding events are likely to be functionally important we further cross-referenced the data with published histone modification ChIP-seq data (Bogdanovic et al., 2012). The 5DE and 5PE1 peaks overlapped significant H3K27ac and H3K4me1 peaks (Table S3), suggesting these are likely to be functionally active enhancers. Tbx16 ChIP-qPCR confirmed the validity of the 5DE and 5PE1 ChIP-seq peaks (Fig. 6C). These putative enhancers are likely to regulate *myf5*, the promoter of which has a H3K4me3 mark, rather than the adjacent *myf6* gene, which is not expressed at 80% epiboly and does not have a H3K4me3 mark. Comparison of the sequences under each peak to genomic regions adjacent to the *myf5* gene in other fish species revealed significant conservation (Figs 6A,B). Of particular note was the conservation of 5DE in medaka and stickleback, while 5PE1 showed notable conservation in medaka and fugu (Figs 6A,B). Thus, ChIP-seq peaks corresponding to histone marks indicative of enhancer activity suggest evolutionarily conserved mechanisms of *myf5* regulation in fish.

**Fig. 6.**
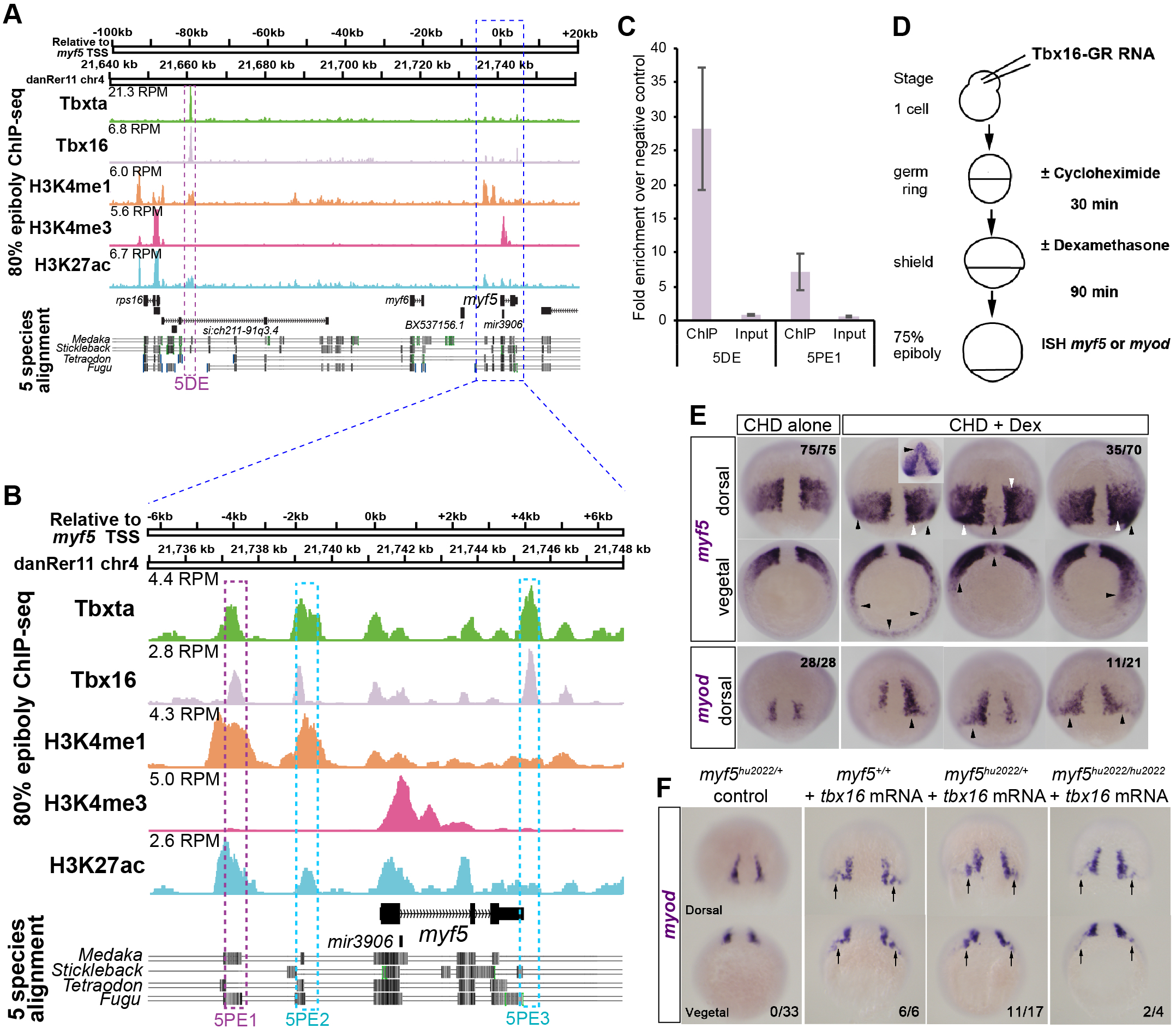
Myf5 is a direct transcriptional target of Tbx16. **A,B.** Chromatin immunoprecipitation followed by sequencing (ChIP-seq) on wild-type (wt) embryos at 75-85% epiboly reveals endogenous Tbx16 and Tbxta binding events within 120 kb flanking the *myf5* transcriptional start site (TSS). RPM – ChIP-seq peak height in reads per million reads. H3K4me3 indicates TSSs. H3K4me1 indicates putative enhancers. H3K27ac indicates active enhancers. Known transcripts with exons (black) and introns (arrowheads) are indicated. Multiz Alignments & Conservation from UCSC Genome Browser (Haeussler et al., 2019) are shown beneath. Purple boxes indicate validated Tbx16 binding sites. Blue box indicates region expanded in panel B. Cyan boxes indicate other Tbx sites mentioned in text. **C**. ChIP-qPCR validation of Tbx16 peaks on *myf5* distal element (5DE) and proximal element 2 (5PE1). Error bars indicate standard error of the mean for biological triplicate experiments. **D.** Schematic showing how direct injection into wt embryos of mRNA encoding the Tbx16-Gulucocotorticoid Response fusion protein leads to target gene induction in the presence of protein synthesis inhibitor cycloheximide (CHD) induced by nuclear translocation triggered by dexamethasone (DEX). CHD caused ∼5% delay in epiboly sowing it was active. **E.** Wild-type embryos injected with *tbx16-GR* mRNA treated with cycloheximide from germ ring stage and dexamethasone from shield stage. Note that embryos treated with cycloheximide alone exhibit wild type *myf5* expression at 75-80% epiboly, whereas embryos additionally treated with dexamethasone exhibit ectopic *myf5* expression with strong (white arrowheads, comparable to wt pre-adaxial level) and weak (black arrowheads, comparable to wt paraxial level) stain. Three separate CDH+DEX treated embryos are shown. Numbers indicate the fraction of embryos with the expression pattern(s) shown. Inset shows an unusual induction of *myf5* in anterior regions that was not observed with *myod*. **F.** Injection of *tbx16* mRNA (200 pg) into embryos from a *myf5^hu2022/+^* heterozygote incross led to ectopic up-regulation of *myod* mRNA in the dorsal germ ring (arrows) irrespective of genotype. Numbers indicate fraction of embryos showing ectopic mRNA/total analysed. Bars: 100 µm.

We next tested whether Tbx16 is able to positively regulate *myf5* directly. To do this we used a hormone-inducible system to activate Tbx16 in the presence of cycloheximide, followed by *in situ* hybridization (Kolm and Sive, 1995; Martin and Kimelman, 2008). Briefly, mRNA corresponding to Tbx16 fused to the hormone binding domain of glucocorticoid receptor (GR) was overexpressed in wild-type embryos. The resulting protein is held in the cytoplasm until dexamethasone (DEX) stimulates GR nuclear translocation. In the presence of the translation inhibitor cycloheximide (CHD), increased nuclear Tbx16 is expected mosaically to induce only direct targets of Tbx16 (Fig. 6D). Ectopic expression of *myf5* around the germ ring was observed upon Tbx16 activation (Fig. 6E). Moreover, using the same approach, induction of *myod* mRNA indicated that *myod* is also a direct target of Tbx16 (Fig. 6E). *Myod* mRNA up-regulation was, however, restricted to a narrow ectopic domain flanking the base of the notochord, indicating that *myod* expression is under additional Tbx16-independent constraints compared to *myf5* (Fig. 6E). Interestingly, across the set of CHD+DEX-treated embryos, ectopic *myf5* mRNA was induced to a higher level in a similar region to ectopic *myod* mRNA than elsewhere, suggesting that Tbx16 was able to induce two aspects of pre-adaxial character directly in this region of the embryo. To confirm this result, we tested whether Myf5 is required for Tbx16 to induce *myod* expression. When *tbx16* mRNA was injected into *myf5* mutant or heterozygote embryos, ectopic *myod* mRNA was observed flanking the base of the notochord in about 50% of embryos, but appeared more readily induced in wild type siblings (Fig. 6F). Thus, Tbx16 is necessary for MRF expression and can induce both *myf5* and *myod* independently, so long as Fgf signalling is active. In summary, Tbx16 directly induces MRF expression in gastrulating mesoderm and is particularly potent in the pre-adaxial region that normally retains high Tbx16 expression.

### Tbxta is essential for pre-adaxial but not paraxial myogenesis

Whereas the entire paraxial PSM expresses *myf5*, pre-adaxial cells upregulate *myf5* and are the first cells to express *myod*. Tbxta and Tbx16 have similar DNA binding recognition sequences (Garnett et al., 2009; Nelson et al., 2017). Congruently, we find a prominent Tbxta ChIP-seq peak at the 5DE −80 kb site upstream of *myf5*, and minor peaks at the proximal sites (Fig. 6A,B; Table S3). Because of the role of Tbx16 and Tbxta in *myod* expression we also examined the *myod* locus for Tbx protein binding. We found multiple sites occupied by Tbxta and Tbx16 either individually or in combination (Table S3). Notably, only one site (DDE3) displayed strongly significant H3K4me1 and H3K27ac peaks and this was only occupied by Tbxta and not by Tbx16 (Fig. S5 and Table S3). However, an additional site (DDE1) showed significant occupancy by Tbx16 and Tbxta concurrent with H3K4me1 but not H3K27ac (Fig. S5; Table S3). In spite of the absence of a significant H3K27ac mark, this peak may be important to Tbx16 regulation of *myod*. These findings indicate that differential direct binding of Tbxta and Tbx16 may control both *myf5* and *myod* expression at the inception of skeletal myogenesis.

Is Tbxta also required for MRF expression in response to Fgf? Over-expression of Fgf4 in *tbxta* mutants successfully induced *myf5* mRNA and suppressed *aplnrb* mRNA widely in the posterior mesoderm except in a widened dorsal midline region, showing that the introduced Fgf4 was active (Fig. 7A,B). However, *myod* expression was not rescued in *tbxta* mutants in the dorsal pre-adaxial region of Fgf4-injected embryos, or elsewhere around the germ ring (Fig. 7A). Moreover, even an increased dose of 225 pg *fgf4* mRNA/embryo failed to rescue *myod* mRNA in *tbxta* mutants. Importantly, *tbxta* heterozygotes showed significantly less extensive induction of *myod* mRNA in response to Fgf4 than did their wt siblings (p=0.0001 Χ^2^-test; Fig. 7C and Table S4). Therefore, Tbxta is essential for *myod* induction in pre-adaxial cells independent of its role in promoting expression of midline Fgfs.

**Fig. 7.**
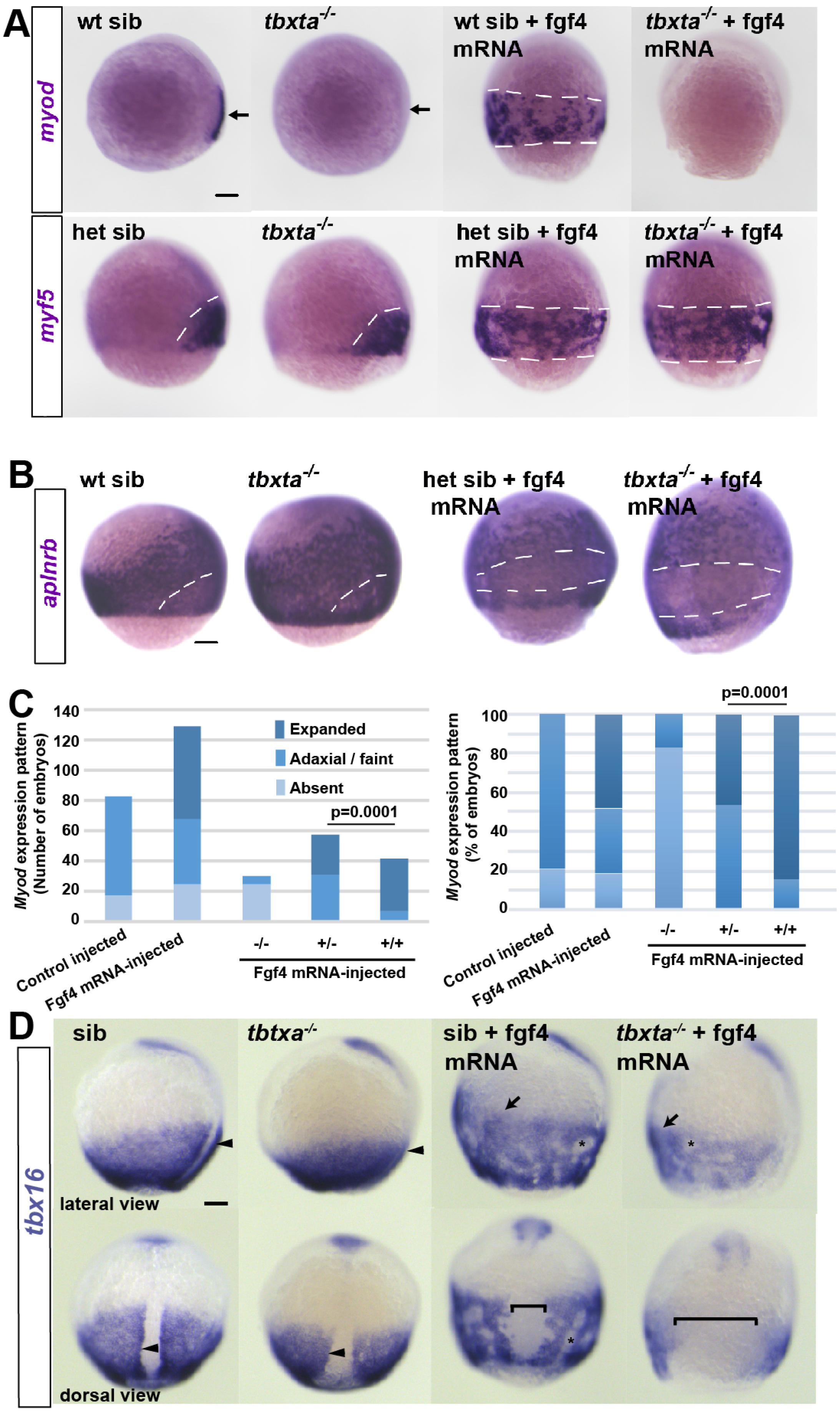
Tbxta is essential for Fgf4-driven induction of *myod* but not *myf5*. Embryos from a *tbxta^+/-^* incross injected with 150 pg *fgf4* mRNA or control. **A.** *Tbtxa^-/-^* mutants lack *myod* mRNA (arrows) but retain *myf5* mRNA in presomitic mesoderm (white dashes). *Fgf4* mRNA injection induced *myf5* and *myod* mRNAs throughout the posterior mesoderm in siblings (white dashes), but failed to induce *myod* mRNA in mutants. **B.** *Fgf4* suppressed *aplnrb* mRNA in posterior mesoderm above the germ ring (white dashes) in both *tbxta^-/-^* mutants and siblings. Individually genotyped embryos are shown in lateral view, dorsal to right (A,B). **C.** Scoring of *myod* mRNA accumulation in response to *Fgf4* mRNA injection into embryos from a *tbxta^+/-^* incross. Expanded: ventral expansion, generally all around germ ring as in panel A. Adaxial/faint: Either wild type pattern or faint version of it in small a small proportion of mutants, which was not significantly altered by Fgf4 mRNA. Left panel shows absolute number of embryos analysed from two experiments to emphasise lack of induction in mutants (raw data in Table S3). Right panel displays data as a percentage of the total to highlight reduced response in heterozygotes compared to wild type (Χ^2^ test). **D.** Adaxial upregulation of *tbx16* mRNA is lost in *tbxta^-/-^* mutant (arrowheads). Fgf4 upregulates *tbx16* mRNA throughout ventral posterior mesoderm (arrows) and causes mesodermal cell aggregation (asterisks). *Tbxta^-/-^* mutants accumulate less *tbx16* mRNA than siblings and have less expression on the dorsal side (brackets). Bars: 100 µm.

Although Fgf4-injection did not radically alter the location of *tbx16* or *tbxta* mRNA (Fig. 5B), we noticed that the higher level of *tbx16* mRNA in adaxial compared to paraxial cells was not obvious in Fgf4-injected wt embryos, with high levels present at all dorsoventral locations, presumably because pre-adaxial character was induced widely in posterior mesoderm (Fig. 5B). Nevertheless, as Fgf4-injection into *tbxta* mutants induced *myf5* but not *myod* mRNA (Fig. 7A), it seems Tbxta is essential to progress from *myf5* to *myod* expression.

Two hypotheses could explain the lack of *myod* expression in Fgf4-injected *tbxta* mutants. First, despite the apparent lack of Tbxta protein in adaxial cells (Ochi et al., 2008; Schulte-Merker et al., 1994a), Tbxta might act directly on *myod*. Alternatively, Fgf-driven Tbxta activity might act indirectly in pre-adaxial cells to upregulate Tbx16 and thereby drive *myod* expression. We therefore examined the ability of Fgf4 to upregulate Tbx16 in *tbxta* mutants. Fgf4 enhanced *tbx16* mRNA throughout the posterior mesoderm in siblings, with the exception of the widened notochordal tissue that contained nuclear Tbxta protein and failed to upregulate MRFs (Figs 7D, 8D). In *tbxta* mutants, Fgf4 also enhanced *tbx16* mRNA in the ventral mesoderm, but a broader dorsal region did not express *tbx16*. Moreover, the level of *tbx16* mRNA appeared lower than in siblings (Fig. 7D). Thus, Fgf4-injected *tbxta* mutants lack both Tbxta and Tbx16 upregulation in pre-adaxial cells. We conclude that Fgf-driven induction of lateral myogenic tissue requires Tbx16, but not Tbxta. In contrast, induction of pre-adaxial character, marked by upregulated *myf5* and *myod* mRNAs, requires both Tbx genes.

**Fig. 8.**
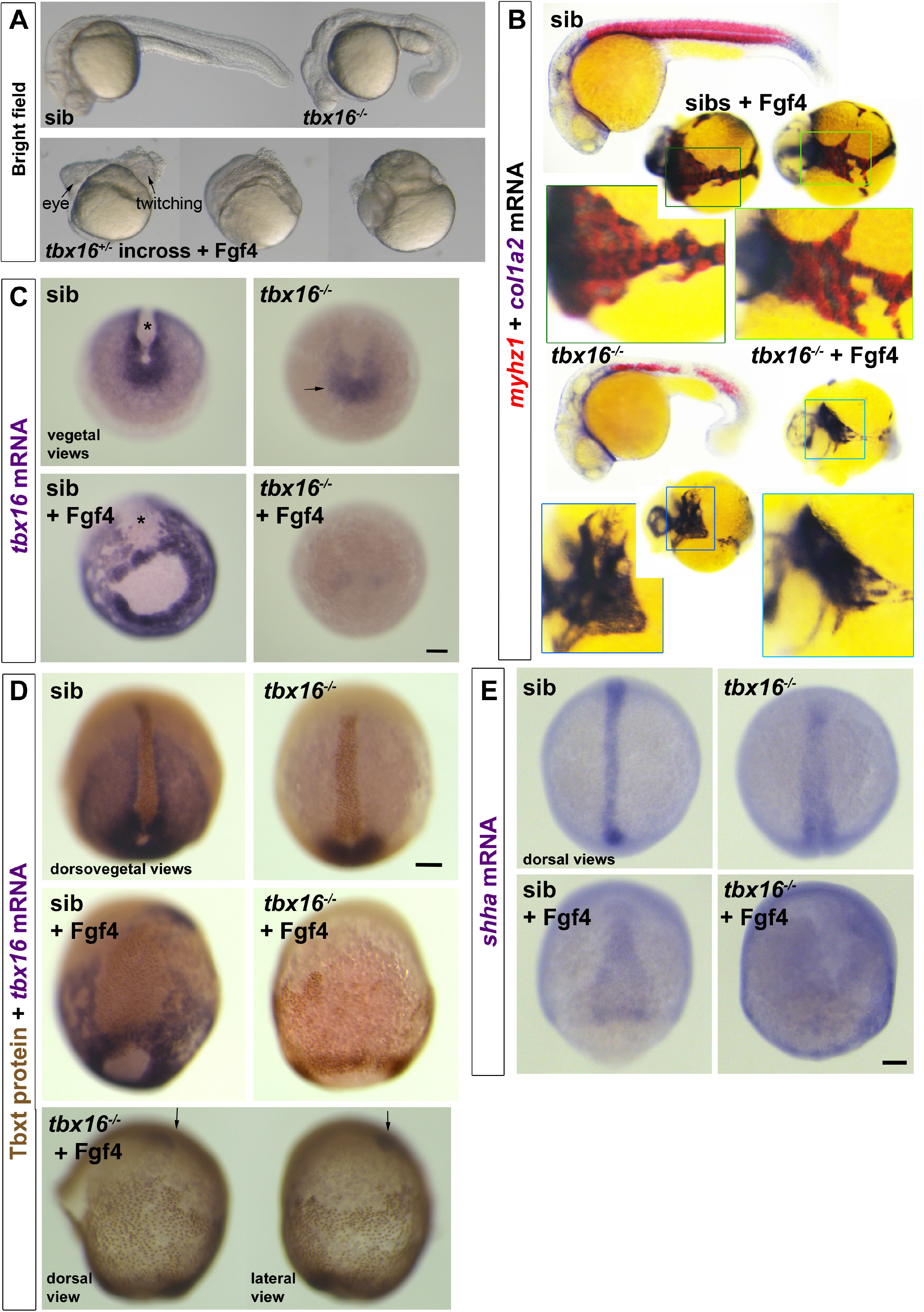
Tbx16 is essential for Fgf4-driven upregulation of *tbx16* and suppression of *tbxta*. Embryos from a *tbx16^+/-^* incross injected with 150 pg *fgf4* mRNA or control. **A,B.** By 24 hpf, Fgf4-injected embryos have disorganized heads and although lacking obvious trunk or tail, some contain twitching muscle (A). In situ mRNA hybridization for *col1a2* for dermomyotome/connective tissue and *myhz1* for skeletal muscle revealed muscle in *fgf4*-injected sibs, but not in *tbx16^-/-^* mutants (B). Boxes are magnified to show the alternating pattern of aggregated muscle and connective tissue in Fgf4-injected siblings, but the reduced *col1a2* and absent *myhz1* mRNA in Fgf4-injected mutants. Note the aggregation of posterior mesoderm cells into strands around the yolk. **C.** In situ mRNA hybridisation for *tbx16* mRNA in embryos from a *tbx16^+/-^* incross at around 90% epiboly. Nonsense mediate decay of the mutant transcript is apparent (arrow). Fgf4 RNA injection increases *tbx16* mRNA in paraxial mesoderm, widens dorsal axial notochord domain (asterisk) and causes aggregation of paraxial cells in siblings, but suppresses residual *tbx16* transcript in mutants. **D.** Immunodetection of Tbxt protein and *tbx16* mRNA in Fgf4-injected and control embryos from a *tbx16^+/-^* incross. A Fgf4-injected *tbx16*^-/-^ mutant (bottom) reveals nuclear Tbxt protein in the entire posterior mesoderm. Residual *tbx16* mRNA in the prechordal region (arrows) but absence in posterior mesoderm demonstrates the genotype. **E.** Widespread up-regulation of *shha* mRNA reveals the notochord-like character of posterior mesoderm in Fgf4-injected *tbx16*^-/-^ mutant. Bars: 100 µm.

### Fgf action on Tbx16 suppresses dorsoposterior axial fate

Finally, we examined the wider effect of Fgf signalling when skeletal muscle cannot form in the absence of *tbx16* function. Excess Fgf action in embryos causes gross patterning defects (Kimelman and Kirschner, 1987; Slack et al., 1987). When Fgf4 was over-expressed in *tbx16* heterozygote in-cross lays, some anterior mesodermal tissue formed and head tissues such as eye and brain were apparent, but trunk and tail mesoderm was grossly deficient (Fig. 8A). In siblings over-expressing Fgf4, some muscle was formed and truncated embryos were observed to twitch at 24 hpf. In contrast, no muscle was detected in *tbx16* mutants (Fig. 8B). Moreover, the residual expression of mutant *tbx16* mRNA at 90% epiboly observed in un-injected embryos was lost upon Fgf4 over-expression (Fig. 8C). This suggests that the cells with tailbud character normally accumulating in *tbx16* mutants were missing. Instead, widespread expression of Tbxta protein in nuclei far from the germ ring suggested that the entire posterior (but not anterior) mesoderm had converted to notochord, the most dorsal posterior mesoderm fate (Fig. 8D). Indeed, *shha* mRNA, a marker of notochord, was found to be broadly upregulated around the embryo in *tbx16* mutant embryos injected with Fgf4 mRNA, but not in their siblings (Fig. 8E). The data suggest that Fgf4 drives early involution of all posterior mesoderm precursors, leaving none to form a tailbud. In the presence of Tbx16, Fgf4 also dorsalizes most involuted mesoderm to muscle, whereas, in the absence of Tbx16, Fgf4 converts most of the mesoderm to notochord.

## Discussion

The current work contains four main findings. First, that Tbx16 directly binds and activates *myf5* regulatory elements to initiate skeletal myogenesis. Second, that Fgf signalling acts through Tbx16 to drive the initial myogenic events in the adaxial cell lineage that subsequently require Hh signalling to complete myogenesis. Third, that Tbxta, the dorsal-most/posterior Tbx factor, binds directly to *myod* regulatory elements and also promotes dorsal midline expression of Fgfs that subsequently cooperate to drive dorsal myogenesis through Tbx16. Fourth, that Fgf action through Tbx16 suppresses the dorsoposterior axial fate induced by Tbxta. Overall, Tbx transcription factors provide a crucial link between mesoderm induction and the initiation of myogenesis, which has profound implications for understanding the evolution of vertebrates.

### Tbx genes and myogenesis

Building on previous evidence that Tbx16 up-regulates *myf5* mRNA level (Garnett et al., 2009; Mueller et al., 2010), our findings that Tbx16 protein binds to DNA elements near the *myf5* and *myod* genes, is required for their expression, and can induce them in the absence of protein synthesis show that these MRF genes are direct targets of Tbx16. We also present evidence that *myf5* and *myod* are direct targets of Tbxta, including the presence of several significant Tbxta ChIP-seq peaks and the requirement for Tbxta to transduce Fgf4-mediated signalling into *myod* expression. As MRF gene activity, principally that of *myf5* and *myod*, drives commitment to skeletal myogenesis in vertebrates, our findings place Tbx protein activity at the base of skeletal myogenesis in zebrafish.

Before myogenesis, Tbx16 is required for migration of most trunk PSM cells away from the ‘maturation zone’ immediately after their involution (Griffin and Kimelman, 2002; Row et al., 2011). Our analysis of *aplnrb*-expressing mesodermal cells shows that most anterior (i.e. head) and posterior ventral (i.e. ventral trunk) mesoderm involution and migration occurs normally in both *tbx16* and *tbxta* mutants. Indeed, some PSM is formed in *tbx16* mutants and goes on to make small amounts of muscle (Amacher et al., 2002; Kimmel et al., 1989). PSM formation is more severely lacking in *tbx16;tbxta* or *tbx16;tbx16l* double mutants (Amacher et al., 2002; Griffin et al., 1998; Morrow et al., 2017; Nelson et al., 2017) or after knockdown of Tbxtb in *tbxta* mutant (Martin and Kimelman, 2008). Cooperation of Tbx proteins in PSM formation also occurs in *Xenopus tropicalis* (Gentsch et al., 2013). It is likely, therefore, that all PSM formation and its accompanying *myf5* expression requires Tbx proteins. Our findings also help to explain the observation that *tbx16* mutants have increased pronephric mesoderm (Warga et al., 2013). As Tbx16 is required for direct induction of *myf5* and for *pcdh8, msgn1, mespaa* and *tbx6* expression in PSM (Fior et al., 2012; Goering et al., 2003; Griffin and Kimelman, 2002; Lee et al., 2009; Morrow et al., 2017; Yamamoto et al., 1998), the data support previous proposals (Amacher and Kimmel, 1998; Griffin and Kimelman, 2002) of a role for Tbx16 in promotion of the earliest step in PSM formation, *en route* to myogenesis. These early actions of Tbx16 and Tbxta proteins have previously masked their direct myogenic actions in mutants.

To identify likely MRF enhancers at 80% epiboly, we have largely restricted our focus to robust Tbx16 and Tbxta ChIP-seq peaks that are co-incident with established histone marks indicative of active enhancers, H3K4me1 and H3K27ac. This approach is driven by the accepted knowledge that most transcription factor binding events are unlikely to be functionally important, and the availability of stage-matched ChIP-seq data for these established histone marks (Bogdanovic et al., 2012). However, there is an increasing realization that comprehensive identification of enhancers will require a more complete analysis of further histone acetylation events, as evidenced by the recent discovery of functional enhancers marked by H3K122ac, H3K64ac and/or H4K16ac, many of which lack H3K27ac (Pradeepa, 2017; Pradeepa et al., 2016). It is therefore possible that additional Tbx16 and Tbxta ChIP-seq peaks beyond 5DE, 5PE1 and DDE3 may mark functionally important enhancers regulating *myf5* and *myod*. Of particular note is DDE1, - 31kb upstream of *myod*, which is colocalized with a significant H3K4me1 mark, though not H3K27ac. Given that *myod* expression is restricted to a minor cell population at 80% epiboly, we further note that the probability of detecting significant histone marks specific to this population is low due to averaging of signal across whole embryos. It is therefore possible that the ChIP-seq peak common to Tbx16 and Tbxta at the DDE1 −31kb upstream of *myod* is functionally important. Given that Tbx16 is essential for *myod* expression in pre-adaxial cells and can upregulate *myod* in the absence of protein synthesis, this ChIP-seq peak may represent a key region mediating this direct transcriptional activation. Our data argue that once posterior (i.e. trunk) mesoderm forms, Tbx proteins are still required for MRF expression and normal myogenesis. Hh signalling from notochord acts to maintain adaxial MRF expression in wild type and, if Tbx-driven initiation fails, Hh can initiate *myod* and up-regulate *myf5* expression, thereby driving slow myogenesis (Blagden et al., 1997; Coutelle et al., 2001; Du et al., 1997). When Hh signalling is prevented, the small amount of trunk PSM that forms in *tbx16* mutants activates low level *myf5* but not *myod* mRNA expression, showing that Tbx16 is essential for initial pre-adaxial *myod* transcription. *Tbxta* mutants also fail to initiate pre-adaxial *myod* expression. Conversely, overexpression of Tbx16 directly induces ectopic *myf5* mRNA, but only in mesoderm and preferentially in dorsal mesoderm at somite levels. In contrast, Tbx16-induced ectopic *myod* expression is restricted to a narrower mesodermal region flanking the pre-adaxial cells, likely due to the restricted expression of *smarcd3* in this region (Ochi et al., 2008). Nevertheless, *myod* is induced by Tbx16 in the absence of Myf5, probably through direct binding to regulatory elements in the *myod* locus. It is thus likely that changes in chromatin structure in the *myf5* and *myod* loci that accompany posterior mesoderm formation facilitate Tbx16 access to its binding sites in these MRF genes.

In considering the role of Tbx genes in myogenesis it is essential to distinguish adaxial and paraxial myogenesis, which give rise to different kinds of muscle with distinct timing (Blagden et al., 1997; Devoto et al., 1996). Paraxial PSM cells form fast muscle after somitogenesis. As posterior mesoderm involutes during gastrulation, *myf5* expression is initiated throughout the PSM. Tbx16, but not Tbxta, is required for almost all this low level *myf5* expression. In wild type, *myf5* mRNA then declines as paraxial PSM matures before being up-regulated once more as somite borders form (Coutelle et al., 2001). In contrast, pre-adaxial cells require both Tbxta and Tbx16 function for up-regulation of *myf5* and initiation of *myod* expression, and then commence terminal differentiation within the PSM to generate slow muscle. Thus, distinct Tbx proteins are required for normal adaxial and paraxial myogenesis.

Our data show that Tbx16 is a key direct regulator of MRFs required for myogenesis. The expression of low levels of *myf5* mRNA and formation of small amounts of both fast and slow muscle in the trunk and more in the tail in *tbx16* mutants are likely driven by Tbx16l (formerly known as Tbx6 and then Tbx6l) (Morrow et al., 2017). Morrow et al (2017) show that tail somite formation is prevented and tail *myod* mRNA diminished in the *tbx16;tbx16l* double mutant, but that *myod* mRNA expression continues at a reduced level throughout the axis at 24 hpf. Our data argue that the residual *myod* mRNA is in adaxially-derived slow muscle induced by Hh signalling.

Other Tbx proteins may directly regulate myogenic initiation in other body regions and species, as many Tbx proteins bind similar DNA motifs (Papaioannou, 2014). For example, Tbx1, Tbx5a and Tbx20 are required in anterior mesoderm for patterning of cranial and cardiac muscles (Knight et al., 2008; Lu et al., 2017; Valasek et al., 2011). *Tbx4/5a* gene function is required for limb muscle patterning, at least partially non-cell autonomously (Don et al., 2016; Hasson et al., 2010; Valasek et al., 2011). Tbx6 suppresses myogenesis in anterior PSM indirectly through Mesp-b (Windner et al., 2015), but direct actions of Tbx6 could also regulate myogenesis. The increasing complexity of Tbx gene diversity during vertebrate evolution may have permitted muscle diversification.

### Fgf and myogenesis

Tbx16 is required for Fgf signalling to induce *myf5* (Fig. 5A). In its absence, Fgf drives all posterior mesoderm to a notochord-like fate (Fig. 8D,E), probably via activation of Tbxta. It has long been known that Fgf signalling is required for expression of *myf5* in the paraxial PSM of the tailbud (Groves et al., 2005). Here we show Fgf is also required for the earliest *myf5* expression in invaginating trunk mesoderm and for the initiation of *myf5* and *myod* expression in pre-adaxial cells destined to form the slow muscle of anterior somites. Our data also extend the evidence that this MRF expression is subsequently maintained, as the shield/tailbud-derived sources of Fgf recede from the adaxial cells, by Hh signalling from the maturing notochord (Coutelle et al., 2001; Osborn et al., 2011). The finding that the latter Fgf action is restricted to trunk (as opposed to tail) somites is consistent with a) the dwindling level of Fgf mRNAs in the chordoneural hinge as tailbud outgrowth slows becoming insufficient to promote MRF expression and b) the unresolved issue of Hh-independent initiation of MRF expression in the anteriormost somites of murine *smoothened* mutants (Zhang et al., 2001). We suggest this MRF expression is Fgf-triggered in mouse, as in zebrafish.

Tbxta is also required for normal Fgf signalling from the midline to promote pre-adaxial myogenesis. One likely reason is the loss of *fgf4, fgf6a*, *fgf8a, fgf8b* and possibly *fgf3* expression in the dorsal midline chordoneural hinge region in *tbxta* mutants (Fig. 3B)(Draper et al., 2003). When Fgf4 or Fgf6a is artificially over-expressed in *tbxta* mutants, *myf5* is readily induced. This suggests that Tbxta protein is not required cell autonomously to drive *myf5* expression, even though our data reveal that Tbxta binds in vivo to similar regions of the *myf5* locus to those bound by Tbx16. In marked contrast, Tbxta is essential for over-expressed Fgf to induce *myod*. Interestingly, ChIP shows Tbxta binding sites upstream of *myod* that are preferentially bound by Tbtxa, suggesting that Tbxta binding to one or more of these sites directly activates *myod* in response to Fgf. We note, however, that Tbxta protein is immunodetectable in notochord and germ ring cells but barely in *myod*-expressing pre-adaxial cells (Hammerschmidt and Nüsslein-Volhard, 1993; Odenthal et al., 1996; Schulte-Merker et al., 1994a). Nevertheless, as pre-adaxial cells have recently expressed Tbxta during their involution, it is possible that functionally significant protein could remain bound at the *myod* locus. Transient persistence of Tbxta is consistent with the downregulation of *myod* mRNA in more anterior adaxial cells in the absence of Hh signals. An alternative hypothesis of indirect regulation of *myod* by Fgf through another transcription factor, such as Tbx16, is also tenable, and is supported by the reduced upregulation of *tbx16* observed in *tbxta* mutants injected with Fgf4 (Fig. 7D). We show that Myf5 is not an indirect mediator of Fgf signalling on *myod*, despite its Fgf sensitivity.

Our data argue strongly that Fgf signalling not only promotes *tbx16* expression, but also enhances the activity of the Tbx16 protein. The MRF inducing activity of Tbx16 is suppressed by inhibition of Fgf signalling (Fig. 5C,D). Bearing in mind the existence of Tbx16l and Tbxta, this result is consistent with the finding that *tbxta* or *tbx16* mutation sensitizes embryos to Fgf inhibition (Griffin and Kimelman, 2003). As Tbx16 overexpression can expand PSM fates and reverse the effect of partial Fgf inhibition, a primary role of Fgf signalling is to cooperate with Tbx16 to drive expression of its target genes, including *myf5*. This understanding provides mechanistic insight into how the effects of Fgf on gastrulation movements and histogenesis are separated, as originally proposed (Amaya et al., 1993). Interestingly, Tbx16 over-expression rescues *myf5* mRNA preferentially on the dorsal side of the embryo, suggesting that BMP and/or other signals continue to suppress PSM fates ventrally, and thus that Tbx16 does not act by simply suppressing the inhibitory effect of such signals.

We find that in the absence of Tbxta, Fgf acts to induce Tbx16 in ventral regions (Fig. 7D). The data are fully consistent with a positive feedback loop between Tbxta and Fgf signalling to maintain tailbud character and notochord formation. Additionally, our findings suggest that Tbx16 acts in competition with Tbxta to suppress tailbud/notochord fate and Fgf expression and to promote paraxial fates. Ochi et al. (2008) have suggested Tbxta and Fgf work through Smarcd3 (Baf60c) to permit pre-adaxial cell formation. It is striking that the requirement for both Tbxta and Tbx16 proteins to generate adaxial cells located between the axial and paraxial regions triggers the first step of diversification of muscle to slow and fast contractile character.

### Evolution of vertebrates

From an evolutionary perspective, our findings yield a number of important insights. As efficient motility is a key feature of animals, and efficient sarcomeric muscle is found throughout triploblasts, it is likely that mesodermal striated muscle existed in the common ancestor of deuterostomes and protostomes. It is widely thought that chordates evolved from an early deuterostome consisting of a segmented ‘branchial basket’, possibly attached to a propulsive segmented pre-anal trunk. There is consensus that subsequent appearance of neural crest, notochord and a post-anal tail were significant evolutionary steps for chordates (Gee, 2018). Already in cephalochordates at least two kinds of striated muscle had evolved in anterior somites (Devoto et al., 2006; Lacalli, 2002). Our evidence that initiation of both slow and fast myogenesis in the most anterior trunk is driven by Fgf/Tbx signalling indicates that a major function of this early mesodermal inducer was induction of trunk striated myogenesis, which may constitute an ancestral chordate character. Once Hh is expressed in maturing midline tissues, it triggers terminal differentiation of muscle precursors into functional muscle through a positive feedback loop (Coutelle et al., 2001; Osborn et al., 2011). Parallel diversification of neural tube cells, also regulated by Hh (Placzek and Briscoe, 2018), may have generated matching motoneural and muscle fibre populations that enhanced organismal motility. With the evolution of a *tbxta*-dependent tailbud destined to make the post-anal tail, our data suggest that weakening Fgf signalling continued to induce *myf5* expression and paraxial mesoderm character through *tbx16*, but was insufficient to induce adaxial myogenesis. The presence of Hh and Tbxta plus Tbx16, however, ensure that adaxial slow muscle is initiated in the zebrafish tail.

In the anterior somites of amniotes, as in zebrafish, Hh signalling maintains, rather than initiates *myf5* expression (Zhang et al., 2001). In more posterior somites of zebrafish, Xenopus and amniote, Hh signalling drives *myf5* initiation (Borycki et al., 1999; Grimaldi et al., 2004). Compared to zebrafish, however, murine *Myf5* induction is further delayed until after somitogenesis, when Gli3 repressive signals in PSM have diminished (McDermott et al., 2005). In mouse, Brachyury/Tbxt is required for myogenesis and binds 20 kb downstream of *Myod*, but does not obviously control its expression (Lolas et al., 2014). No clear orthologue of *tbx16* exists in mammals, although it clusters by sequence with *Tbx6* genes. In mammals, Tbx6 suppresses neurogenesis in posterior paraxial mesoderm, suggesting that additional mechanisms have evolved that suppress early pre-somitic *Myf5* expression (Chapman and Papaioannou, 1998). Indeed, possible low level *Myf5* expression in PSM has long been a source of controversy (George-Weinstein et al., 1996; Gerhart et al., 2004). Thus, there has been diversification in how these Tbx genes regulate somitic myogenesis.

Our data suggest that regions required for *myf5* expression are located ∼80 kb upstream of the transcription start site. A BAC transgenic encompassing 80 kb upstream and 76 kb downstream of *myf5* has been shown to drive GFP expression in muscle (Chen et al., 2007). However, analysis of shorter constructs has been confounded by cloning artefacts (Chen et al., 2001; Chen et al., 2007), leaving understanding of elements driving specific aspects of zebrafish *myf5* expression unclear. Murine *Myf5* is also regulated by many distant elements (Buckingham and Rigby, 2014). Similarly, we observe Tbx binding peaks far upstream of zebrafish *myod*. Upstream elements are known to initiate murine *Myod* expression in some embryonic regions, but whether these elements drive the earliest myotomal regulation of *Myod* is unknown (Chen and Goldhamer, 2004). The extent to which similar transcription factors act through similar binding elements to initiate MRF expression and myogenesis across vertebrates remains to be determined.

The ancestral situation seems clearer. In amphioxus, *Tbx6/16* is expressed in tailbud and PSM (Belgacem et al., 2011). In *Ciona*, knockdown of Tbx6b/c/d leads to reduced *MyoD* expression, loss of muscle and paralysis (Imai et al., 2006). In *Xenopus*, both Tbx6 and VegT are implicated in early myogenesis (Callery et al., 2010; Fukuda et al., 2010; Tazumi et al., 2010), although some mechanisms may differ from those in zebrafish (Maguire et al., 2012). By adding our zebrafish findings, we show that in all major chordate groups Tbx-dependent gene regulation is central to skeletal myogenesis. The conserved involvement, yet divergent detail, of how *Tbx16*, *Tbx6* and *Tbxt* genes regulate somitic myogenic diversity along the body axis are consistent with selective pressures on these duplicated Tbx gene families playing a key role in the diversification of myogenesis in the vertebrate trunk and tail, characters that gave chordates their predatory advantage.

## Materials and methods

### Zebrafish lines and maintenance

Mutant lines *fgf8a^ti282a^* (Reifers et al., 1998), *noto^n1^* (Talbot et al., 1995), *smo^b641^* (Barresi et al., 2000), *tbxta^b195^* and *tbx16^b104^* (Griffin et al., 1998) are likely nulls and were maintained on King’s wild type background. Staging and husbandry were as described previously (Westerfield, 2000). All experiments were performed under licences awarded under the UK Animal (Scientific Procedures) Act 1986 and subsequent modifications.

### In situ mRNA hybridization and immunohistochemistry

In situ mRNA hybridization for *myf5* and *myod* was as described previously (Hinits et al., 2009). Additional probes were *fgf3* (Maroon et al., 2002), *fgf4* (IMAGE: 6790533), *fgf6*a (Thisse and Thisse, 2005), *fgf8a* (Reifers et al., 1998), *tbxta* (Schulte-Merker et al., 1994b) and *tbx16* (Griffin et al., 1998; Ruvinsky et al., 1998). Anti-Ntla immunostaining was performed after in situ hybridization using rabbit anti-Ntla (Schulte-Merker et al., 1992, 1:2000) and Horse anti-rabbit IgG-HRP conjugated (Vector Laboratories, Inc.)

### Embryo Manipulations

Embryos were injected with MOs (GeneTools LLC) as indicated in Table S2 to *fgf3*, *fgf4*, *fgf6a*, *fgf8a*, *tbxta* (Feldman and Stemple, 2001) or *tbx16* (Bisgrove et al., 2005). Controls were vehicle or irrelevant mismatch MO. Cyclopamine (100 μM in embryo medium), SU5402 (at indicted concentration in embryo medium) and vehicle control were added at 30% epiboly to embryos whose chorions had been punctured with a 30G hypodermic needle. A PCR product of *fgf4* (IMAGE: 6790533) was cloned (primers in Table S2) into the SacI/SalI sites of pßUT3 to make mRNA for over-expression. 100-220 pg *fgf4* mRNA (made with messageMachine), 50 pg *fgf6a* mRNA, 150 pg *tbx16* mRNA (Griffin et al., 1998) or 150 pg *tbx16*-GR mRNA (Jahangiri et al., 2012) were injected into 1-cell-stage embryos. For hormone-inducible Tbx16 activation, embryos were treated with 10 μg/ml final concentration of cycloheximide two hours prior to collection at 75-80% epiboly. After 30 min, 20 μM dexamethasone was added for the remaining 1.5 h.

### Chromatin immunoprecipitation and sequencing (ChIP-seq) and ChIP-qPCR

ChIP-seq data was analysed as reported previously (Nelson et al., 2017). ChIP-qPCR experiments were performed as previously reported (Jahangiri et al., 2012) using the primers in Table S2.

## Acknowledgements

We are grateful to all members of the Hughes lab for advice and to Bruno Correia da Silva and his staff for care of the fish. SMH is an MRC Scientist with MRC Programme Grant (G1001029 and MR/N021231/1) support. This work was also supported by grants from the British Heart Foundation to YH and SMH (PG PG/14/12/30664) and to FCW (PG/13/19/30059).

---

**Table S1.**
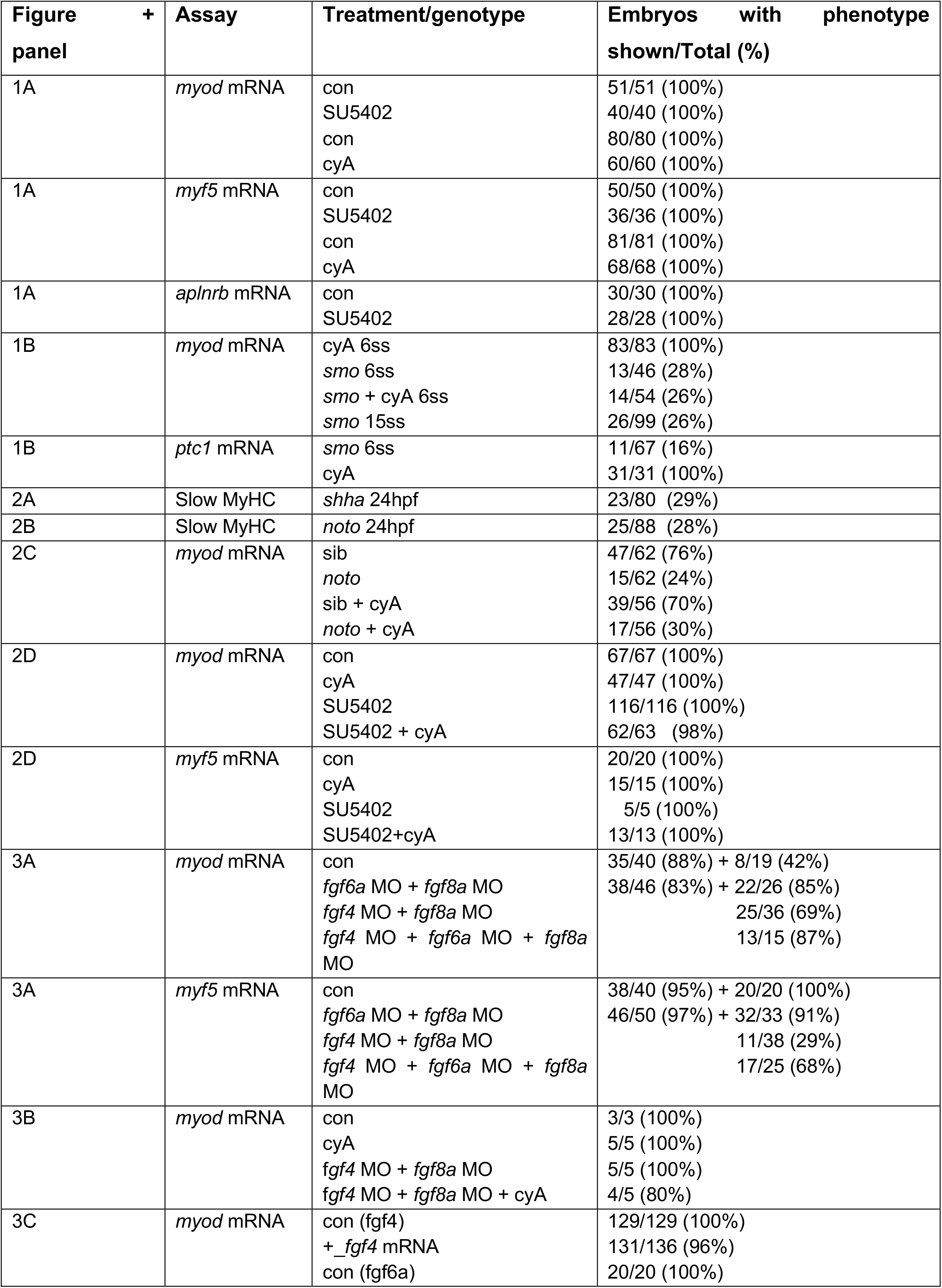
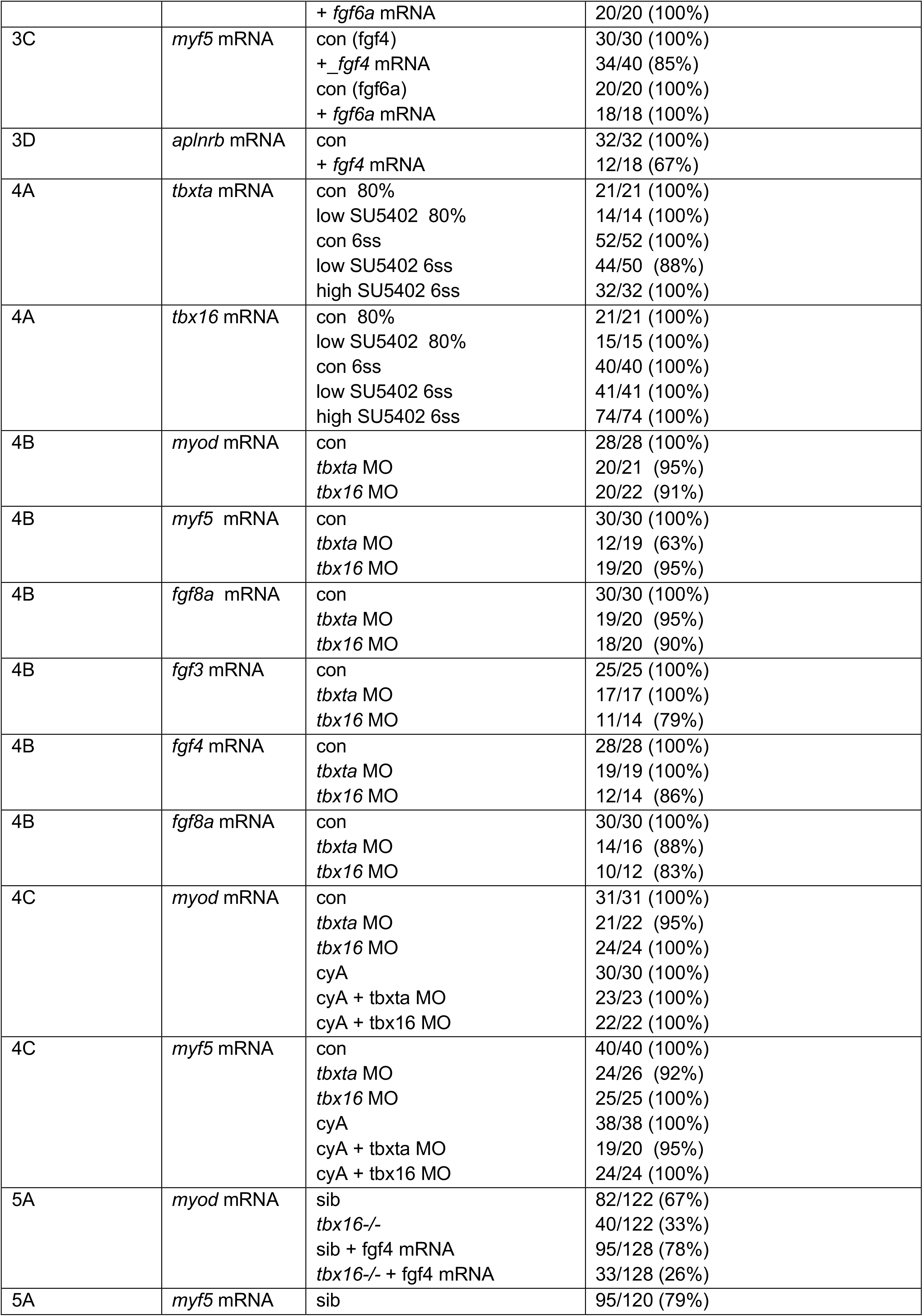
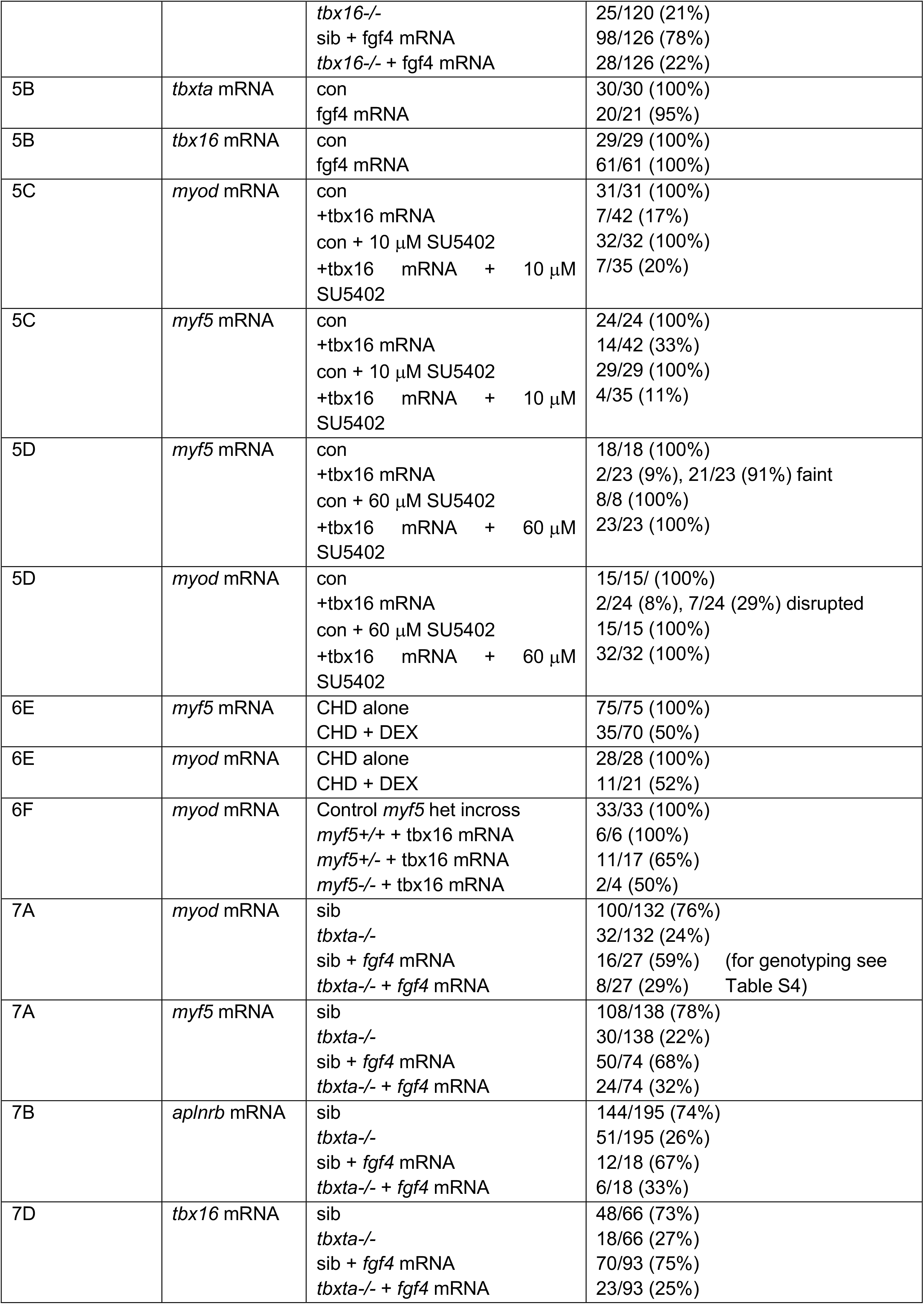
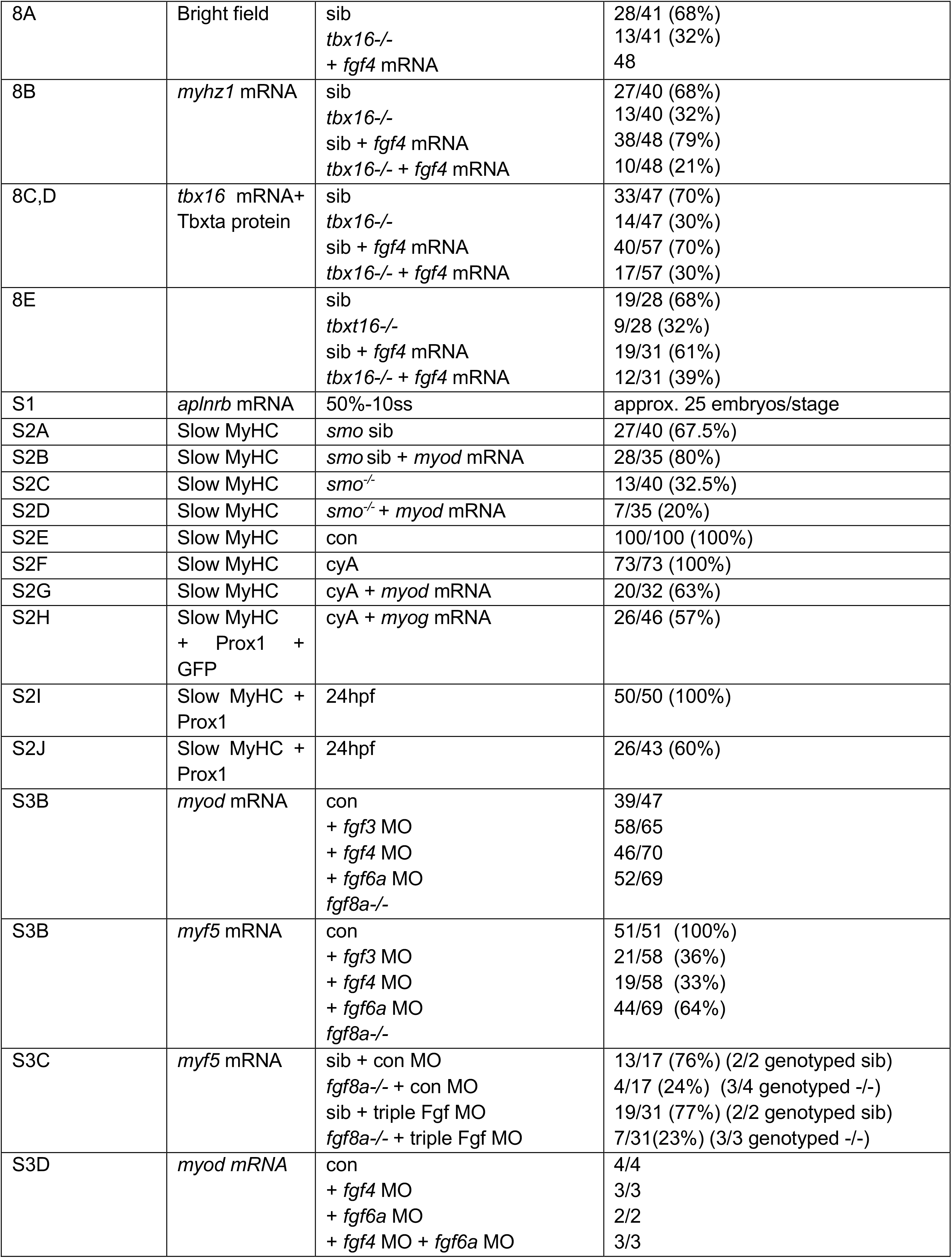
Quantitation of data in Figures.

| Figure panel | + | Assay | Treatment/genotype | Embryos with phenotype shown/Total (%) |
| --- | --- | --- | --- | --- |
| 1A |  | *myod* mRNA | con<br>SU5402<br>con<br>cyA | 51/51 (100%)<br>40/40 (100%)<br>80/80 (100%)<br>60/60 (100%) |
| 1A |  | *myf5* mRNA | con<br>SU5402<br>con<br>cyA | 50/50 (100%)<br>36/36 (100%)<br>81/81 (100%)<br>68/68 (100%) |
| 1A |  | *aplnrb* mRNA | con<br>SU5402 | 30/30 (100%)<br>28/28 (100%) |
| 1B |  | *myod* mRNA | cyA 6ss<br>*smo* 6ss<br>*smo* + cyA 6ss<br>*smo* 15ss | 83/83 (100%)<br>13/46 (28%)<br>14/54 (26%)<br>26/99 (26%) |
| 1B |  | *ptc1* mRNA | *smo* 6ss<br>cyA | 11/67 (16%)<br>31/31 (100%) |
| 2A |  | Slow MyHC | *shha* 24hpf | 23/80 (29%) |
| 2B |  | Slow MyHC | *noto* 24hpf | 25/88 (28%) |
| 2C |  | *myod* mRNA | sib<br>*noto*<br>sib + cyA<br>*noto* + cyA | 47/62 (76%)<br>15/62 (24%)<br>39/56 (70%)<br>17/56 (30%) |
| 2D |  | *myod* mRNA | con<br>cyA<br>SU5402<br>SU5402 + cyA | 67/67 (100%)<br>47/47 (100%)<br>116/116 (100%)<br>62/63 (98%) |
| 2D |  | *myf5* mRNA | con<br>cyA<br>SU5402<br>SU5402+cyA | 20/20 (100%)<br>15/15 (100%)<br>5/5 (100%)<br>13/13 (100%) |
| 3A |  | *myod* mRNA | con<br>*fgf6a* MO + *fgf8a* MO<br>*fgf4* MO + *fgf8a* MO<br>*fgf4* MO + *fgf6a* MO + *fgf8a* MO | 35/40 (88%) + 8/19 (42%)<br>38/46 (83%) + 22/26 (85%)<br>25/36 (69%)<br>13/15 (87%) |
| 3A |  | *myf5* mRNA | con<br>*fgf6a* MO + *fgf8a* MO<br>*fgf4* MO + *fgf8a* MO<br>*fgf4* MO + *fgf6a* MO + *fgf8a* MO | 38/40 (95%) + 20/20 (100%)<br>46/50 (92%) + 32/33 (97%)<br>11/38 (29%)<br>17/25 (68%) |
| 3B |  | *myod* mRNA | con<br>cyA<br>*fgf4* MO + *fgf8a* MO<br>*fgf4* MO + *fgf8a* MO + cyA | 3/3 (100%)<br>5/5 (100%)<br>5/5 (100%)<br>4/5 (80%) |
| 3C |  | *myod* mRNA | con (*fgf4*)<br>+ *fgf4* mRNA<br>con (*fgf6a*) | 129/129 (100%)<br>131/136 (96%)<br>20/20 (100%) |

|  |  |  |  |
| --- | --- | --- | --- |
|  |  | + *fgf6a* mRNA | 20/20 (100%) |
| 3C | *myf5* mRNA | con (*fgf4*)<br>+ *_fgf4* mRNA<br>con (*fgf6a*)<br>+ *fgf6a* mRNA | 30/30 (100%)<br>34/40 (85%)<br>20/20 (100%)<br>18/18 (100%) |
| 3D | *aplnrb* mRNA | con<br>+ *fgf4* mRNA | 32/32 (100%)<br>12/18 (67%) |
| 4A | *tbxta* mRNA | con 80%<br>low SU5402 80%<br>con 6ss<br>low SU5402 6ss<br>high SU5402 6ss | 21/21 (100%)<br>14/14 (100%)<br>52/52 (100%)<br>44/50 (88%)<br>32/32 (100%) |
| 4A | *tbx16* mRNA | con 80%<br>low SU5402 80%<br>con 6ss<br>low SU5402 6ss<br>high SU5402 6ss | 21/21 (100%)<br>15/15 (100%)<br>40/40 (100%)<br>41/41 (100%)<br>74/74 (100%) |
| 4B | *myod* mRNA | con<br>*tbxta* MO<br>*tbx16* MO | 28/28 (100%)<br>20/21 (95%)<br>20/22 (91%) |
| 4B | *myf5* mRNA | con<br>*tbxta* MO<br>*tbx16* MO | 30/30 (100%)<br>12/19 (63%)<br>19/20 (95%) |
| 4B | *fgf8a* mRNA | con<br>*tbxta* MO<br>*tbx16* MO | 30/30 (100%)<br>19/20 (95%)<br>18/20 (90%) |
| 4B | *fgf3* mRNA | con<br>*tbxta* MO<br>*tbx16* MO | 25/25 (100%)<br>17/17 (100%)<br>11/14 (79%) |
| 4B | *fgf4* mRNA | con<br>*tbxta* MO<br>*tbx16* MO | 28/28 (100%)<br>19/19 (100%)<br>12/14 (86%) |
| 4B | *fgf8a* mRNA | con<br>*tbxta* MO<br>*tbx16* MO | 30/30 (100%)<br>14/16 (88%)<br>10/12 (83%) |
| 4C | *myod* mRNA | con<br>*tbxta* MO<br>*tbx16* MO<br>cyA<br>cyA + *tbxta* MO<br>cyA + *tbx16* MO | 31/31 (100%)<br>21/22 (95%)<br>24/24 (100%)<br>30/30 (100%)<br>23/23 (100%)<br>22/22 (100%) |
| 4C | *myf5* mRNA | con<br>*tbxta* MO<br>*tbx16* MO<br>cyA<br>cyA + *tbxta* MO<br>cyA + *tbx16* MO | 40/40 (100%)<br>24/26 (92%)<br>25/25 (100%)<br>38/38 (100%)<br>19/20 (95%)<br>24/24 (100%) |
| 5A | *myod* mRNA | sib<br>*tbx16*<sup>-/-</sup><br>sib + *fgf4* mRNA<br>*tbx16*<sup>-/-</sup> + *fgf4* mRNA | 82/122 (67%)<br>40/122 (33%)<br>95/128 (78%)<br>33/128 (26%) |
| 5A | *myf5* mRNA | sib | 95/120 (79%) |
|  |  | *tbx16*<sup>-/-</sup><br>sib + *fgf4* mRNA<br>*tbx16*<sup>-/-</sup> + *fgf4* mRNA | 25/120 (21%)<br>98/126 (78%)<br>28/126 (22%) |
| 5B | *tbxta* mRNA | con<br>*fgf4* mRNA | 30/30 (100%)<br>20/21 (95%) |
| 5B | *tbx16* mRNA | con<br>*fgf4* mRNA | 29/29 (100%)<br>61/61 (100%) |
| 5C | *myod* mRNA | con<br>+ *tbx16* mRNA<br>con + 10 μM SU5402<br>+ *tbx16* mRNA + 10 μM SU5402 | 31/31 (100%)<br>7/42 (17%)<br>32/32 (100%)<br>7/35 (20%) |
| 5C | *myf5* mRNA | con<br>+ *tbx16* mRNA<br>con + 10 μM SU5402<br>+ *tbx16* mRNA + 10 μM SU5402 | 24/24 (100%)<br>14/42 (33%)<br>29/29 (100%)<br>4/35 (11%) |
| 5D | *myf5* mRNA | con<br>+ *tbx16* mRNA<br>con + 60 μM SU5402<br>+ *tbx16* mRNA + 60 μM SU5402 | 18/18 (100%)<br>2/23 (9%), 21/23 (91%) faint<br>8/8 (100%)<br>23/23 (100%) |
| 5D | *myod* mRNA | con<br>+ *tbx16* mRNA<br>con + 60 μM SU5402<br>+ *tbx16* mRNA + 60 μM SU5402 | 15/15/ (100%)<br>2/24 (8%), 7/24 (29%) disrupted<br>15/15 (100%)<br>32/32 (100%) |
| 6E | *myf5* mRNA | CHD alone<br>CHD + DEX | 75/75 (100%)<br>35/70 (50%) |
| 6E | *myod* mRNA | CHD alone<br>CHD + DEX | 28/28 (100%)<br>11/21 (52%) |
| 6F | *myod* mRNA | Control *myf5* het incross<br>*myf5*<sup>+/+</sup> + *tbx16* mRNA<br>*myf5*<sup>+/-</sup> + *tbx16* mRNA<br>*myf5*<sup>-/-</sup> + *tbx16* mRNA | 33/33 (100%)<br>6/6 (100%)<br>11/17 (65%)<br>2/4 (50%) |
| 7A | *myod* mRNA | sib<br>*tbxta*<sup>-/-</sup><br>sib + *fgf4* mRNA<br>*tbxta*<sup>-/-</sup> + *fgf4* mRNA | 100/132 (76%)<br>32/132 (24%)<br>16/27 (59%)<br>8/27 (29%) (for genotyping see Table S4) |
| 7A | *myf5* mRNA | sib<br>*tbxta*<sup>-/-</sup><br>sib + *fgf4* mRNA<br>*tbxta*<sup>-/-</sup> + *fgf4* mRNA | 108/138 (78%)<br>30/138 (22%)<br>50/74 (68%)<br>24/74 (32%) |
| 7B | *aplnrb* mRNA | sib<br>*tbxta*<sup>-/-</sup><br>sib + *fgf4* mRNA<br>*tbxta*<sup>-/-</sup> + *fgf4* mRNA | 144/195 (74%)<br>51/195 (26%)<br>12/18 (67%)<br>6/18 (33%) |
| 7D | *tbx16* mRNA | sib<br>*tbxta*<sup>-/-</sup><br>sib + *fgf4* mRNA<br>*tbxta*<sup>-/-</sup> + *fgf4* mRNA | 48/66 (73%)<br>18/66 (27%)<br>70/93 (75%)<br>23/93 (25%) |
| 8A | Bright field | sib<br>*tbx16*<sup>-/-</sup><br>+ *fgf4* mRNA | 28/41 (68%)<br>13/41 (32%)<br>48 |
| 8B | *myhz1* mRNA | sib<br>*tbx16*<sup>-/-</sup><br>sib + *fgf4* mRNA<br>*tbx16*<sup>-/-</sup> + *fgf4* mRNA | 27/40 (68%)<br>13/40 (32%)<br>38/48 (79%)<br>10/48 (21%) |
| 8C,D | *tbx16* mRNA+<br>Tbxta protein | sib<br>*tbx16*<sup>-/-</sup><br>sib + *fgf4* mRNA<br>*tbx16*<sup>-/-</sup> + *fgf4* mRNA | 33/47 (70%)<br>14/47 (30%)<br>40/57 (70%)<br>17/57 (30%) |
| 8E |  | sib<br>*tbxt16*<sup>-/-</sup><br>sib + *fgf4* mRNA<br>*tbx16*<sup>-/-</sup> + *fgf4* mRNA | 19/28 (68%)<br>9/28 (32%)<br>19/31 (61%)<br>12/31 (39%) |
| S1 | *aplnrb* mRNA | 50%-10ss | approx. 25 embryos/stage |
| S2A | Slow MyHC | *smo* sib | 27/40 (67.5%) |
| S2B | Slow MyHC | *smo* sib + *myod* mRNA | 28/35 (80%) |
| S2C | Slow MyHC | *smo*<sup>-/-</sup> | 13/40 (32.5%) |
| S2D | Slow MyHC | *smo*<sup>-/-</sup> + *myod* mRNA | 7/35 (20%) |
| S2E | Slow MyHC | con | 100/100 (100%) |
| S2F | Slow MyHC | cyA | 73/73 (100%) |
| S2G | Slow MyHC | cyA + *myod* mRNA | 20/32 (63%) |
| S2H | Slow MyHC<br>+ Prox1 +<br>GFP | cyA + *myog* mRNA | 26/46 (57%) |
| S2I | Slow MyHC +<br>Prox1 | 24hpf | 50/50 (100%) |
| S2J | Slow MyHC +<br>Prox1 | 24hpf | 26/43 (60%) |
| S3B | *myod* mRNA | con<br>+ *fgf3* MO<br>+ *fgf4* MO<br>+ *fgf6a* MO<br>*fgf8a*<sup>-/-</sup> | 39/47<br>58/65<br>46/70<br>52/69 |
| S3B | *myf5* mRNA | con<br>+ *fgf3* MO<br>+ *fgf4* MO<br>+ *fgf6a* MO<br>*fgf8a*<sup>-/-</sup> | 51/51 (100%)<br>21/58 (36%)<br>19/58 (33%)<br>44/69 (64%) |
| S3C | *myf5* mRNA | sib + con MO<br>*fgf8a*<sup>-/-</sup> + con MO<br>sib + triple Fgf MO<br>*fgf8a*<sup>-/-</sup> + triple Fgf MO | 13/17 (76%) (2/2 genotyped sib)<br>4/17 (24%) (3/4 genotyped -/-)<br>19/31 (77%) (2/2 genotyped sib)<br>7/31(23%) (3/3 genotyped -/-) |
| S3D | *myod* mRNA | con<br>+ *fgf4* MO<br>+ *fgf6a* MO<br>+ *fgf4* MO + *fgf6a* MO | 4/4<br>3/3<br>2/2<br>3/3 |

**Table S2.**
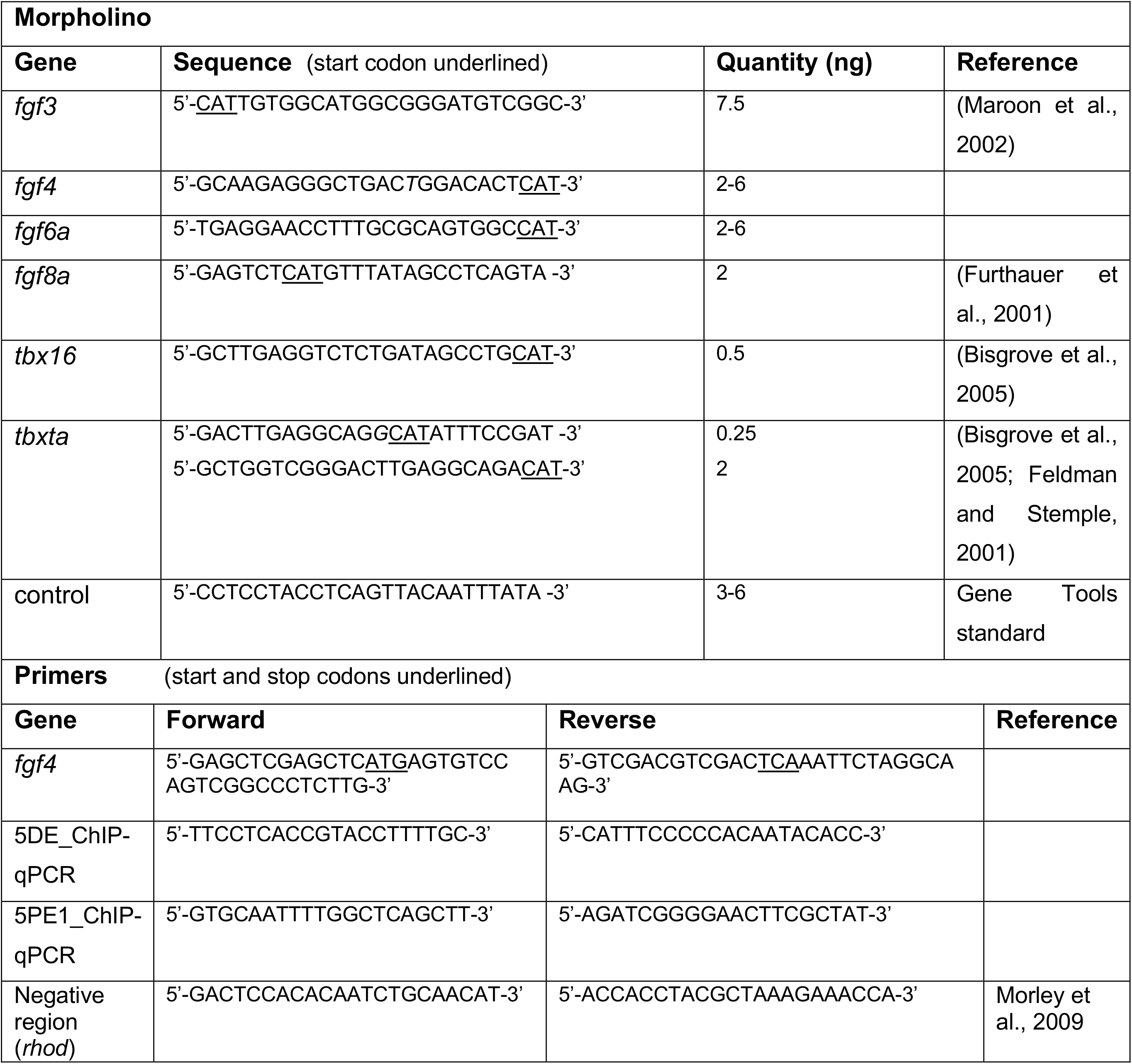
Sequences of morpholinos and primers.

**Table S3.**
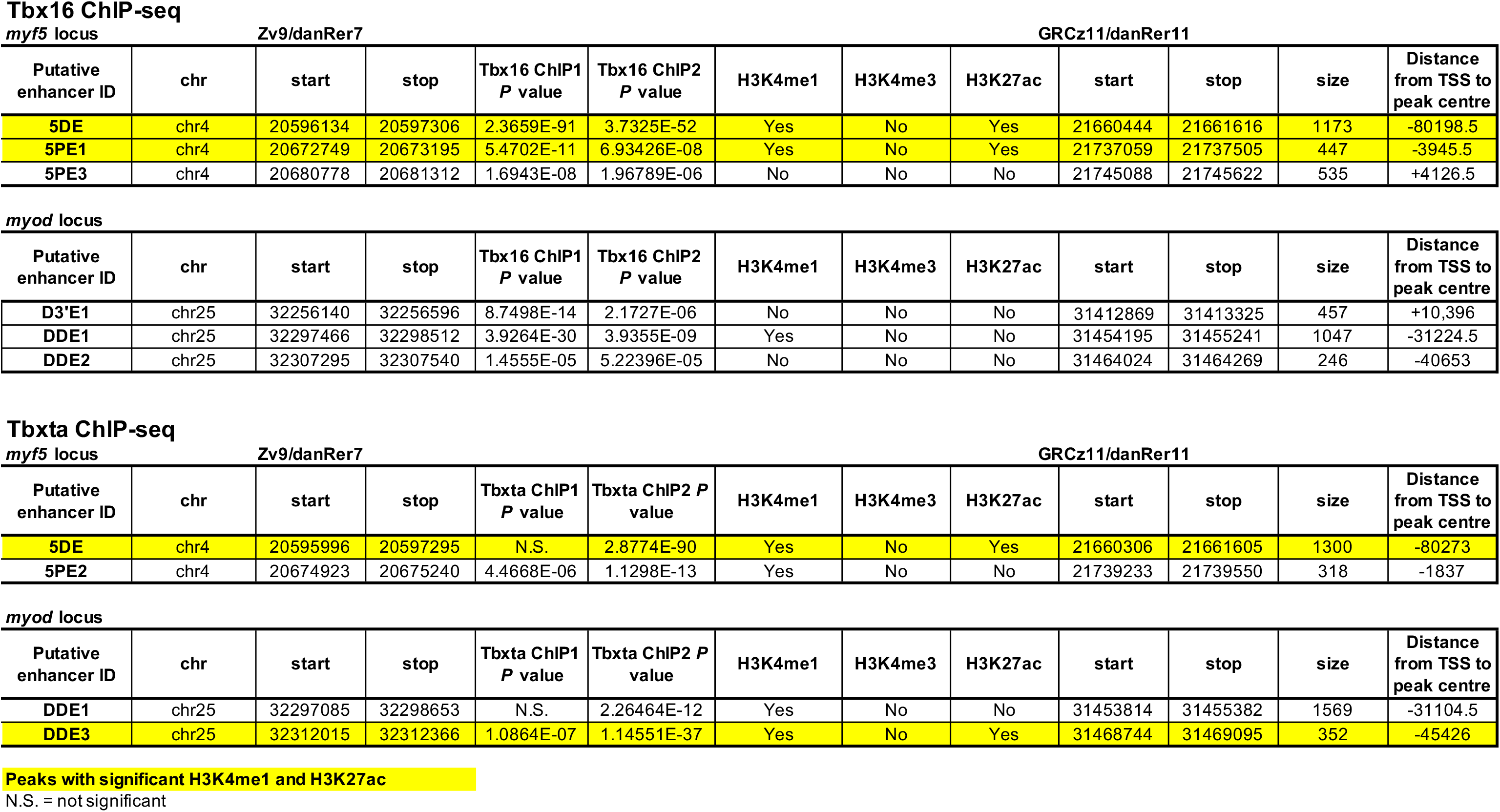
Location and histone modifications of Tbx16 and Tbxta ChIP-seq peaks on *myf5* and *myod* loci.

**Table S4.** Tbxta dosage controls 1220 response of *myod* to Fgf.

| Fgf4 mRNA (pg) | Genotype | Number genotyped [%]¶ | *Myod* mRNA expression pattern in genotyped embryos (%) |  |  |
| --- | --- | --- | --- | --- | --- |
|  |  |  | Absent | Adaxial only | Expanded ventrally |
| 0 | -/- | 4* | 4 (100%)§ | 0 (0%) | 0 (0%) |
|  | +/- | 3* | 0 (0%) | 3 (100%) | 0 (0%) |
|  | +/+ | 2* | 0 (0%) | 2 (100%) | 0 (0%) |
|  | Total | 9 | 8 ((15%))∞ | 45 ((85%)) | 0 ((0%)) |
| 100 | -/- | 8 [30%] | 7 (88%) | 1 (12%) | 0 (0%) |
|  | +/- | 14 [52%] | 0 (0%) | 11 (79%) | 3 (21%) |
|  | +/+ | 5 [19%] | 0 (0%) | 0 (0%) | 5 (100%) |
|  | Total | 27 | 7 (26%) | 12 (44%) | 8 (30%) |
| 0 | -/- | 5* | 5 (100%) | 0 (0%) | 0 (0%) |
|  | +/- | 2* | 0 (0%) | 2 (100%) | 0 (0%) |
|  | +/+ | 3* | 0 (0%) | 3 (100%) | 0 (0%) |
|  | Total | 10 | 9 ((30%)) | 21 ((70%)) | 0 ((0%)) |
| 100 | -/- | 1 [6%] | 1 (100%) | 0 (0%) | 0 (0%) |
|  | +/- | 8 [50%] | 0 (0%) | 3 (37%) | 5 (63%) |
|  | +/+ | 7 [44%] | 0 (0%) | 1 (14%) | 6 (86%) |
|  | Total | 16 | 1 (6%) | 4 (25%) | 11 (69%) |
| 150 | -/- | 6 [25%] | 5 (100%) | 1 (17%) | 0 (0%) |
|  | +/- | 9 [38%] | 0 (0%) | 6 (67%) | 3 (33%) |
|  | +/+ | 9 [38%] | 0 (0%) | 2 (22%) | 7 (78%) |
|  | Total | 24 | 5 (21%) | 9 (38%) | 10 (42%) |
| 225 | -/- | 15 [24%] | 12 (80%) | 3 (20%) | 0 (0%) |
|  | +/- | 26 [42%] | 0 (0%) | 11 (42%) | 15 (58%) |
|  | +/+ | 21 [34%] | 0 (0%) | 4 (19%) | 17 (81%) |
|  | Total | 62 | 12 (19%) | 18 (29%) | 32 (52%) |
| **Summary** |  |  |  |  |  |
| 0 | -/- | 9* | 9 (100%) | 0 (0%) | 0 (0%) |
|  | +/- | 5* | 0 (0%) | 5 (100%) | 0 (0%) |
|  | +/+ | 5* | 0 (0%) | 5 (100%) | 0 (0%) |
|  | Total | 19 | 17 ((20%)) | 66 ((80%)) | 0 ((0%)) |
| 100-225 | -/- | 30 [23%] | 25 (88%) | 5 (12%) | 0 (0%) |
|  | +/- | 57 [44%] | 0 (0%) | 31 (54%) | 26 (46%) |
|  | +/+ | 42 [33%] | 0 (0%) | 7 (17%) | 35 (83%) |
|  | Total | 129 | 25 (19%) | 43 (33%) | 61 (47%) |

\* Only a subset of control embryos were genotyped (to ensure reproducibility).
§ Percentages in curved brackets represent fraction of embryos of indicated genotype showing listed *myod* expression pattern.
¶ Percentages in square brackets represent fraction of embryos in sample with each genotype.
∞ Percentages in double brackets represent fraction of embryos in sample showing listed *myod* expression pattern.

**Fig. S1.**
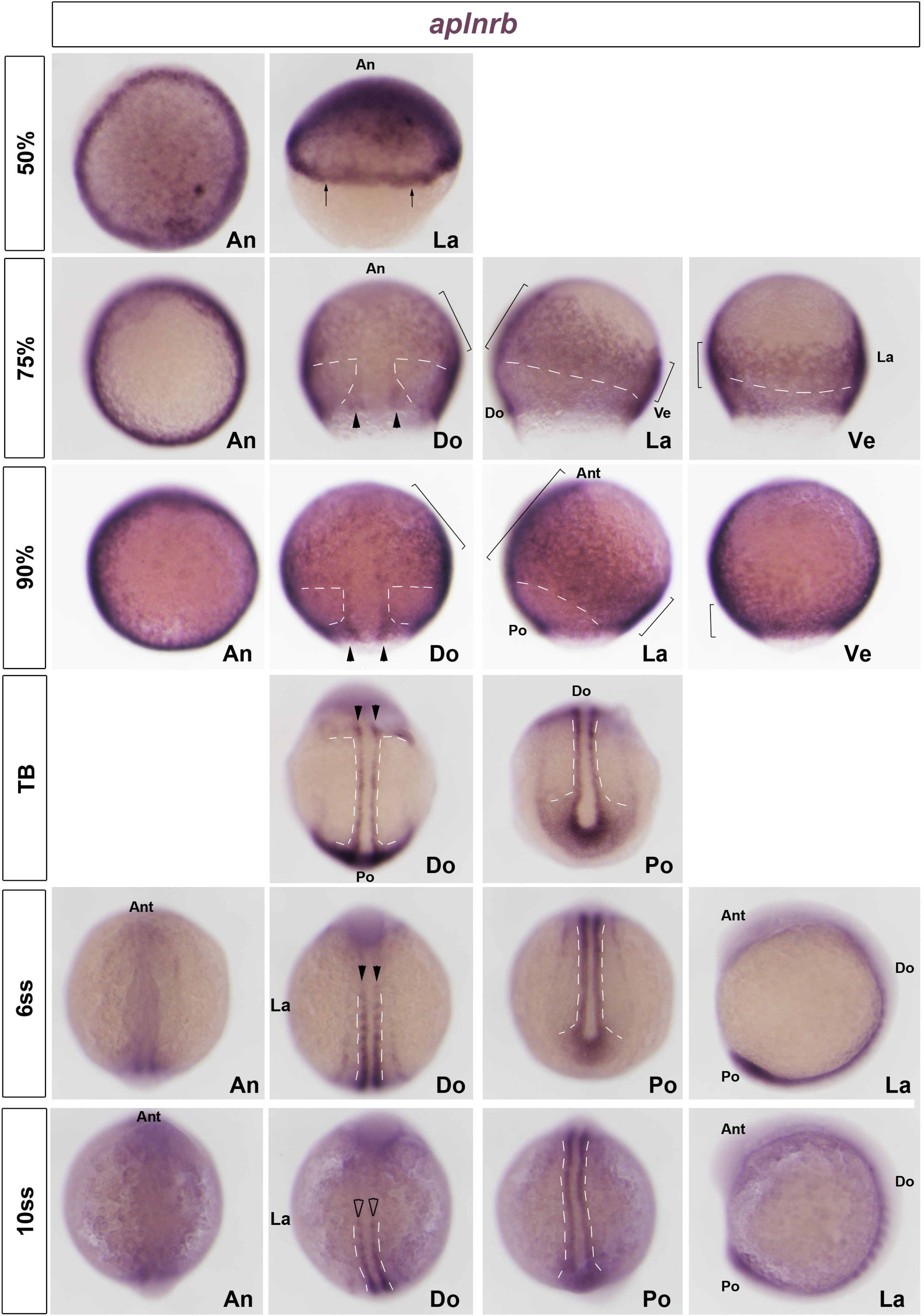
Expression of *aplnrb* mRNA during zebrafish axis formation. Wholemount in situ mRNA hybridization of apelin receptor b *(aplnrb)* mRNA in zebrafish embryos at the indicated stages, shown in animal (An), lateral (La), dorsal (Do, animal to top), ventral (Ve, animal to top) and posterior (Po, dorsal to top) views. Ant = anterior. Note significant expression in early germ ring (arrows), future cranial mesoderm (large and small brackets highlight comparable regions of expression) and adaxial cells (arrowheads). Expression is lacking in paraxial mesoderm (white dashes) that expresses *myf5* and later *myod* mRNAs (see Fig. 1C).

**Fig. S2.**
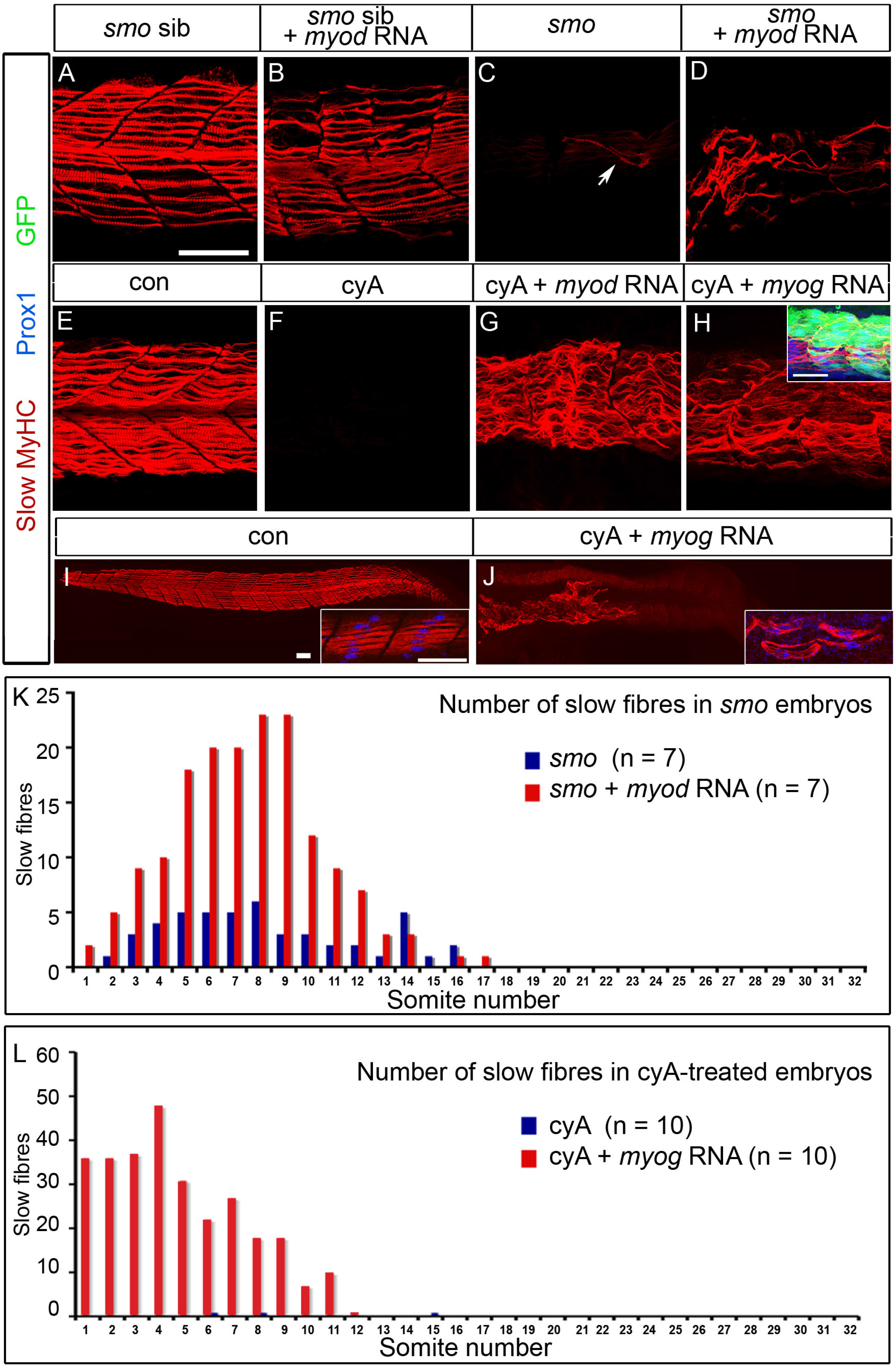
MRF over-expression rescues trunk slow myogenesis. Confocal stacks showing immunodetection of slow fibres with Slow MyHC in *smo* mutant (identified by lack of tail circulation), *smo* sibling, un-injected control or cyA-treated embryos injected with *myod* or *myog* RNA. All embryos orientated anterior to left dorsal up showing 2-3 trunk somites (A-H) or entire trunk and tail (I,J). **A,B.** *Myod* RNA-injected *smo* siblings have slow muscle with disrupted somite morphology**. C,D.** Rare slow fibres present in the trunk region of *smo* mutants (arrow) are more common after *myod* RNA injection. **E,F.** CyA-treatment prevents slow fibre formation. Presence of maternal Smo protein may account for the greater number of residual slow fibres in *smo* mutant compared to cyA-treated embryos. **G,H.** *Myod* or *myog* RNA injection rescues slow fibre formation in cyA-treated embryos. Inset in H shows co-expression of slow MyHC, Prox1 and GFP in a cyA-treated embryo after injection of *myog-IRES-GFP* RNA. **I,J.** *Myog* RNA rescues slow fibres in trunk but not tail. Insets show co-expression of Prox1 and slow MyHC in short confocal stacks. **K.** Slow fibres were counted at 24 hpf in each somite of seven control *smo* mutants and seven *smo* mutants injected at 1 cell stage with *myod* RNA. **L.** Slow fibres were counted at 24 hpf in each somite of ten control uninjected and ten embryos injected at 1 cell stage with *myog* RNA that were each subsequently treated with cyA from 30% epiboly. Bars: 50 µm.

**Fig. S3.**
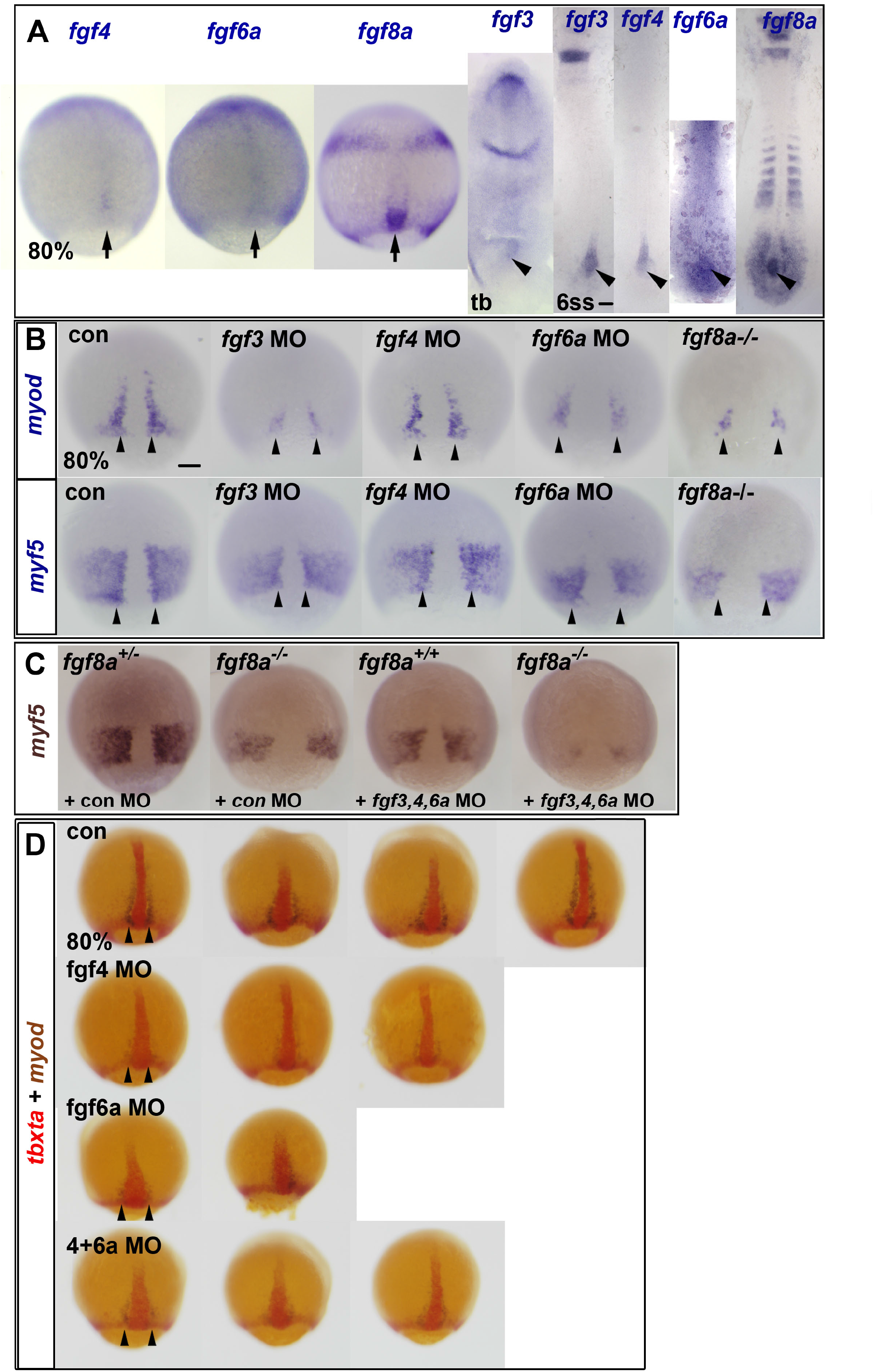
Dorsal fgf expression and requirement for adaxial myogenesis. In situ mRNA hybridization for fgfs in wild type embryos at 80% epiboly, tailbud (tb) and 6ss (A), for *myod* and *myf5* in control and fgf MO-injected and *fgf8a^-/-^* embryos at 80% (B,C) and for *tbxta* (red) and *myod* (blue/brown) (D). **A.** *fgf8a, fgf4*, *fgf6a* and *fgf3* transcripts appear successively in wild type embryos in the dorsal midline (arrows) and CNH (arrowheads). **B.** *Myod* and *myf5* mRNAs in *fgf3* MO, *fgf4* MO and *fgf6a* MO wild type embryos and in sequence-genotyped *fgf8a^-/-^* embryos at 80% epiboly (upper rows, con and single MOs from a representative experiment). Note that siblings of the *fgf8a* mutants had similar MRF expression. Arrowheads indicate nascent adaxial cells. **C.** *Myf5* mRNA in sibling embryos from an incross of heterozygous *fgf8a^+/^*^-^ fish injected with 6 ng control MO or 2 ng each of *fgf3*, *fgf4* and *fgf6a* MO. Note the successively stronger reduction in signal as more fgf function is removed. **D.** Rows showing replicate fgf MO-injected embryos had reduced accumulation of *myod* mRNA in pre-adaxial cells of compared to control (con) (arrowheads). Note widening of notochord in *fgf6a* morphants. Bars: 100 µm.

**Fig. S4.**
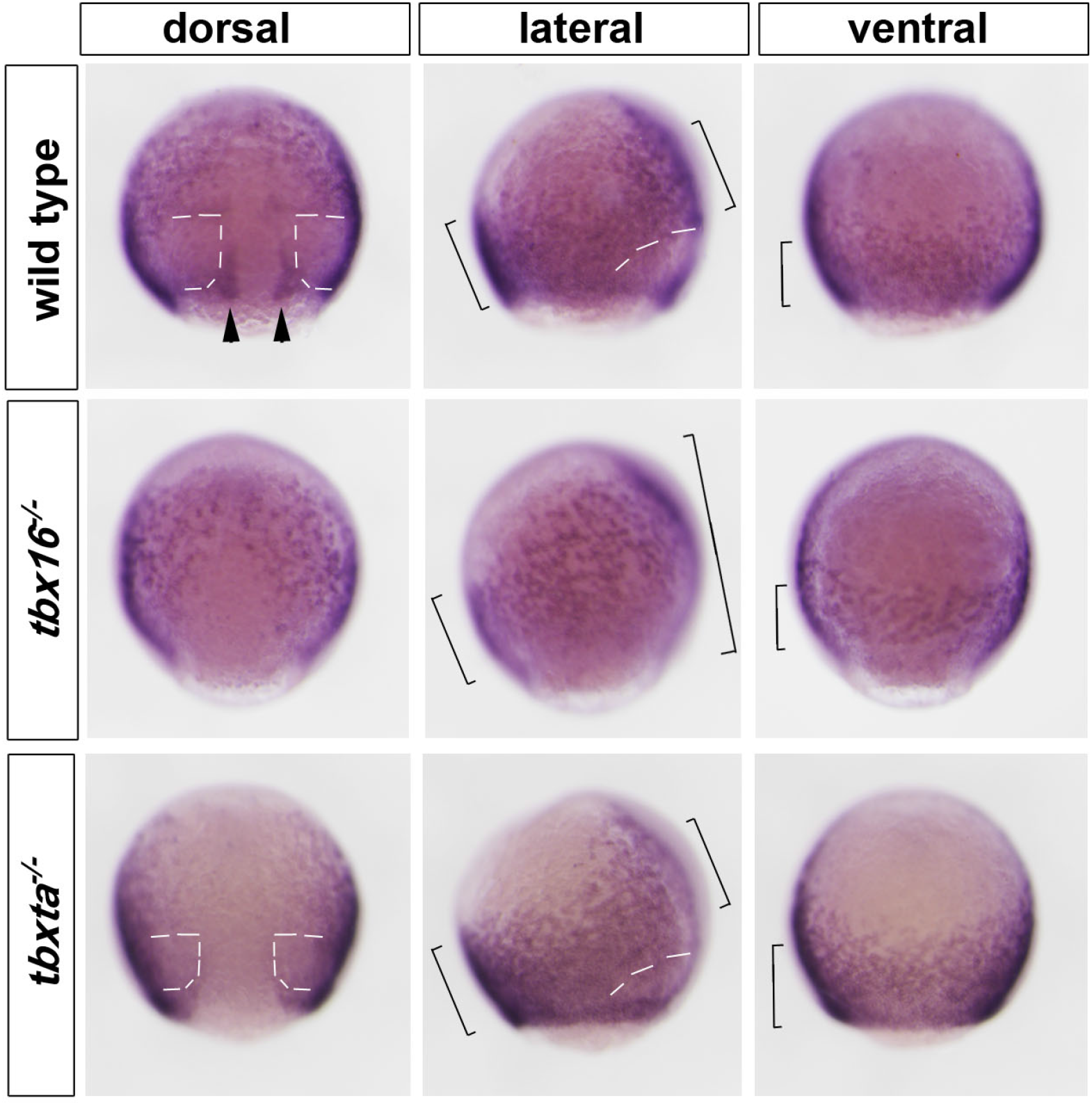
*Aplnrb* mRNA changes in Tbx mutants. In situ mRNA hybridization for *aplnrb* in wild type sibling and *tbx16* mutant and *tbxta* mutant embryos at 80% epiboly. Single embryos are shown from dorsal, left lateral and ventral views. Labelling (brackets) is in a band of anterior mesoderm. Note the unlabelled region in wild type and *tbxta* mutant that is missing in *tbx16* mutant (white dashes). Adaxial *aplnrb* mRNA up-regulation (arrowheads) is lacking in mutants.

**Fig. S5.**
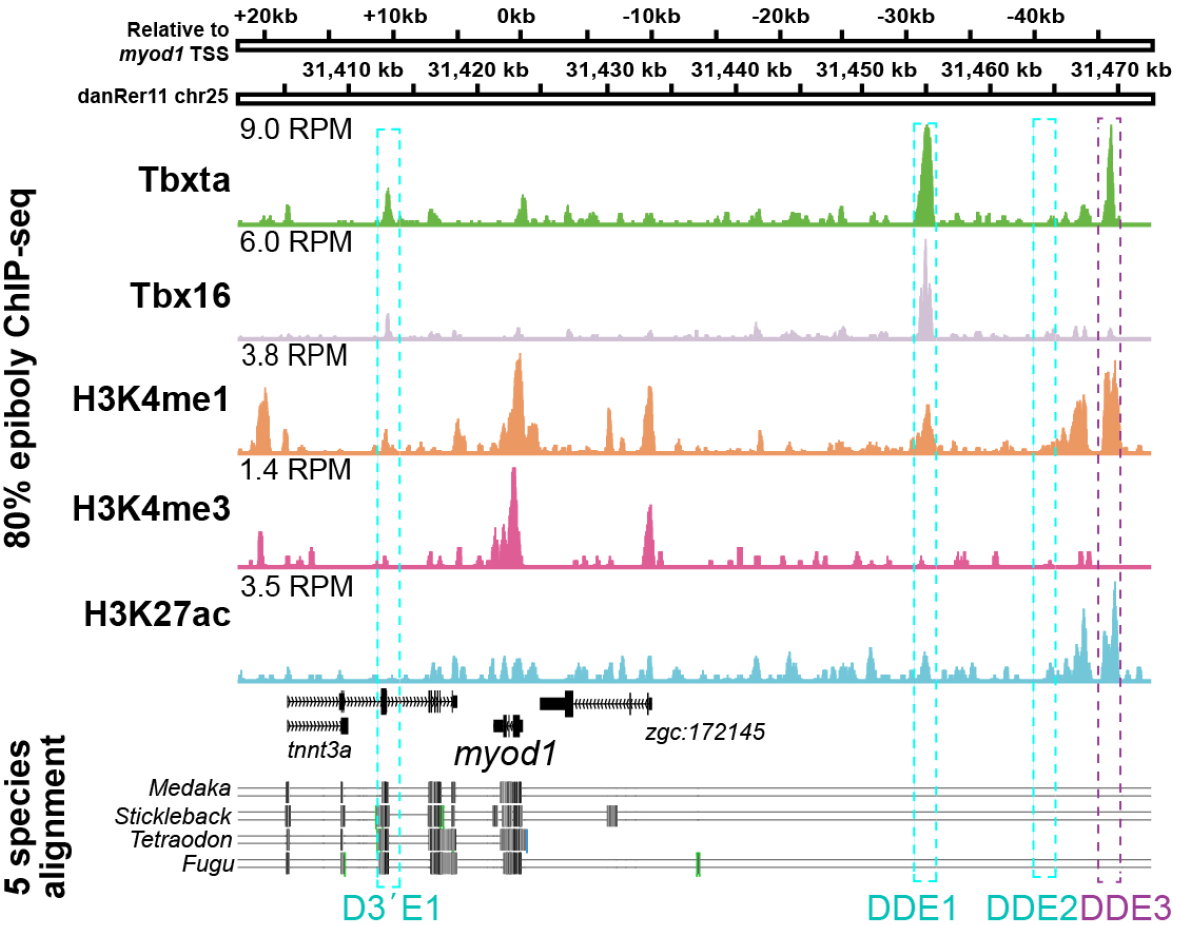
ChIP-seq analysis of *myod* locus. ChIP-seq on wt embryos at 75-85% epiboly indicates endogenous Tbx16 and Tbxta binding events within 75 kb flanking *myod* TSS. H3K4me3 marks TSSs; H3K4me1 marks enhancers; H3K27ac indicates active enhancers; RPM – ChIP-seq peaks height in reads per million reads. Multiz Alignments & Conservation from UCSC Genome Browser (Haeussler et al., 2019) are shown beneath. Purple boxes indicate significant Tbx binding for Tbx16 and Tbxta (DDE1) and Tbxta alone (DDE3). Cyan boxes indicate of Tbx sites mentioned in text. Significant H3K4me1 marks are present at both DDE1 and DDE3, while only DDE3 has a significant H3K27ac mark.

